# The mitotic spindle mediates nuclear migration through an extremely narrow infection structure of the rice blast fungus *Magnaporthe oryzae*

**DOI:** 10.1101/2021.04.07.438902

**Authors:** Mariel A. Pfeifer, Chang Hyun Khang

## Abstract

The blast fungus, *Magnaporthe oryzae*, causes severe destruction to rice and other crops worldwide. As the fungus infects rice, it develops unique cellular structures, such as an appressorium and a narrow penetration peg, to permit successful invasion of host rice cells. Fundamental knowledge about these cellular structures and how organelles, such as the nucleus, are positioned within them is still emerging. Previous studies show that a single nucleus becomes highly stretched during movement through the narrow penetration peg in an extreme nuclear migration event. Yet, the mechanism permitting this nuclear migration event remains elusive. Here, we investigate the role of the mitotic spindle in mediating nuclear migration through the penetration peg. We find that disruption of spindle function during nuclear migration through the penetration peg prevents development of invasive hyphae and virulence on rice. Furthermore, regulated expression of conserved kinesin motor proteins, MoKin5 and MoKin14, is essential to form and maintain the spindle, as well as, properly nucleate the primary hypha. Overexpression of MoKin5 leads to formation of aberrant microtubule protrusions, which contributes to formation of nuclear fragments within the appressorium and primary hypha. Conversely, overexpression of MoKin14 causes the spindle to collapse leading to the formation of monopolar spindles. These results establish a mechanistic model towards understanding the intricate subcellular dynamics of extreme nuclear migration through the penetration peg, a critical step in the development of rice blast disease.

**Importance:** *Magnaporthe oryzae*, also known as the blast fungus, is a formidable hinderance to global food production, including rice. The destructive fungal pathogen develops highly-specialized cells and structures, such as appressoria and penetration pegs, to permit successful invasion of rice cells. Our understanding of *M. oryzae’s* fundamental biology during host cell invasion and colonization is still developing. For instance, it is not yet known how organelles, such as the nucleus, migrate through the narrow penetration peg. Moreover, few previous studies examine the role of motor proteins in *M. oryzae.* In this study, we determined that the mitotic spindle propels a single nucleus through the penetration peg to permit successful development of fungal hyphae inside the first-invaded rice cell. We also identified two conserved kinesin motor proteins, MoKin5 and MoKin14. Our analyses suggested that MoKin5 and MoKin14 exhibit canonical functions in *M. oryzae* during rice infection. This study addressed long-standing questions in rice blast biology, and our results offer opportunities for future research.

## Introduction

Nuclear migration and proper nuclear positioning are fundamental eukaryotic processes. Disruption of nuclear migration, which can lead to improper nuclear positioning, is linked to developmental defects in lower eukaryotes and disease states in humans and higher eukaryotic organisms (1). Seminal studies of nuclear migration and positioning in fungi revealed that cellular components, such as microtubules (MTs) and motor proteins (i.e., kinesins and dynein) are required for successful nuclear migration (2–4). Studies of nuclear migration in various organisms underscore that mechanisms of nuclear migration can be complex, involving the eloquent coordination of cytoskeletons and various motor proteins within the context of the cell cycle. Mechanisms of nuclear positioning vary in fungi (5). For example, in mature hyphae of the ascomycete *Neurospora crassa*, cytoplasmic bulk flow passively moves nuclei forward (6). Other fungi, like the basidiomycete *Ustilago maydis*, utilize a mitotic nuclear migration event to deliver a newly-divided nucleus to the bud (7). In nuclear migration events that occur during mitosis, the spindle is a key player. Spindles are elaborate cellular machines that ensure genetic information is equally divided between mother and daughter cells. Spindles are comprised of MTs, spindle pole bodies (SPBs), and condensed chromosomes called chromatids, along with motor and other MT-associated proteins.

One powerful framework used to explain the intricacy of spindle formation, as well as elongation and maintenance of the spindle throughout mitosis is the force-balance model (8). The force-balance model establishes that spindles are formed and maintained by motor proteins exerting antagonizing forces upon SPBs. In many fungi and other eukaryotes, these motor proteins are members of the kinesin-5 and kinesin-14 superfamilies. Canonical functions of kinesin-5 and kinesin-14 motor proteins are defined. Kinesin-5 motor proteins walk towards the growing plus-ends of MTs and exert an outward force on SPBs (9). Kinesin-14 motor proteins walk towards the minus-ends of MTs and exert an inward force on SPBs (10). However, not all eukaryotes rely on kinesin-5 and kinesin-14 motor proteins to form and maintain a spindle. For example, kinesin-5 is dispensable in the human pathogenic fungus, *Candida albicans* (11). Within *Drosophila* embryos, dynein provides the antagonistic inward force instead of kinesin-14 (12). While the mitotic roles of kinesin-5 and kinesin-14 are defined during development of a number of model organisms, much less is known about the functions of these proteins in spindle formation and function in diverse biological contexts. For instance, what are the roles of these motor proteins in forming a spindle within eukaryotic pathogens as pathogens infect hosts?

The blast fungus, *Magnaporthe oryzae* (anamorph *Pyricularia oryzae),* is a plant pathogen capable of grievous damage to cereal crops worldwide, including rice, wheat and finger millet (13–16). The *M. oryzae* and rice pathosystem serves as a valuable model towards understanding the nuclear migration dynamics of a pathogen during host infection. Rice blast infection is initiated when conidia of *M. oryzae* attach to rice leaves. Conidia germinate and develop appressoria. Appressoria are highly-melanized infection structures. Within appressoria, huge amounts of turgor pressure accumulate, and cytoskeletons, such as F-actins and septins, rearrange at the appressorial pore to give rise to the first fungal structure to enter the rice cell, the penetration peg (17, 18). The penetration peg is narrow (∼0.7 µm) (19). As the fungus continues to grow inside the first-invaded rice cell, bulbous invasive hyphae (IH) develop. Once the first-invaded rice cell is completely colonized by the fungus, the fungus seeks pit fields, housing plasmodesmata, to continue proliferating within rice cells (20). At plasmodesmata in the first-invaded rice cell, the fungus develops another narrow structure called the IH peg (21). The IH peg serves as a conduit to connect IH within the first-invaded rice cell to IH growing within adjacent rice cells. Eventually disease lesions appear on the surface of rice leaves as the fungus spreads throughout the plant.

The nuclear migration dynamics of *M. oryzae* are best characterized during vegetative hyphal growth and during the early events of rice infection (22). *Magnaporthe oryzae* is mononuclear, i.e., each cell contains a single nucleus. During early rice infection, a single nucleus, referred to here as the mother nucleus, is located within the appressorium (23, 24). The newly formed migrating nucleus, here called the daughter nucleus, endures an extreme nuclear migration event, while the mother nucleus remains within the appressorium. (Fig. 1A). During this extreme nuclear migration event, the mother nucleus with a diameter of ∼ 2 µm begins to divide within the appressorium. Subsequently, the daughter nucleus becomes highly stretched (Fig. 1A, middle panel) as it transits the constricted penetration peg that has a diameter of ∼ 0.7 µm (23). The daughter nucleus then quickly moves to the apical region of the primary hypha located inside the first-invaded rice cell (23, 24). Typically, this process lasts ∼5 minutes, with the daughter nucleus traveling over 20 µm from the appressorium through the penetration peg into the primary hypha (23). Although the general behavior of the mother and daughter nucleus are characterized during this extreme nuclear migration event, the cytoskeletons involved in this process are unknown. Based on studies of subsequent *M. oryzae* infection stages, it is likely the spindle is involved. During development of bulbous IH inside the first-invaded rice cell, the spindle nucleates newly-formed IH during mitotic nuclear migration (25, 26). The spindle also delivers a newly-formed nucleus to IH growing in adjacent rice cells through the narrow IH peg. During movement through the IH peg, the spindle can adopt a striking geometry to facilitate movement of the nucleus (26). Intriguingly, the migrating nucleus with a diameter of ∼2µm becomes highly elongated as it moves through the IH peg with a diameter of ∼0.5, which is akin to the nuclear morphology of the migrating daughter nucleus during movement through the penetration peg at earlier stages of rice infection (25).

**Fig. 1.**
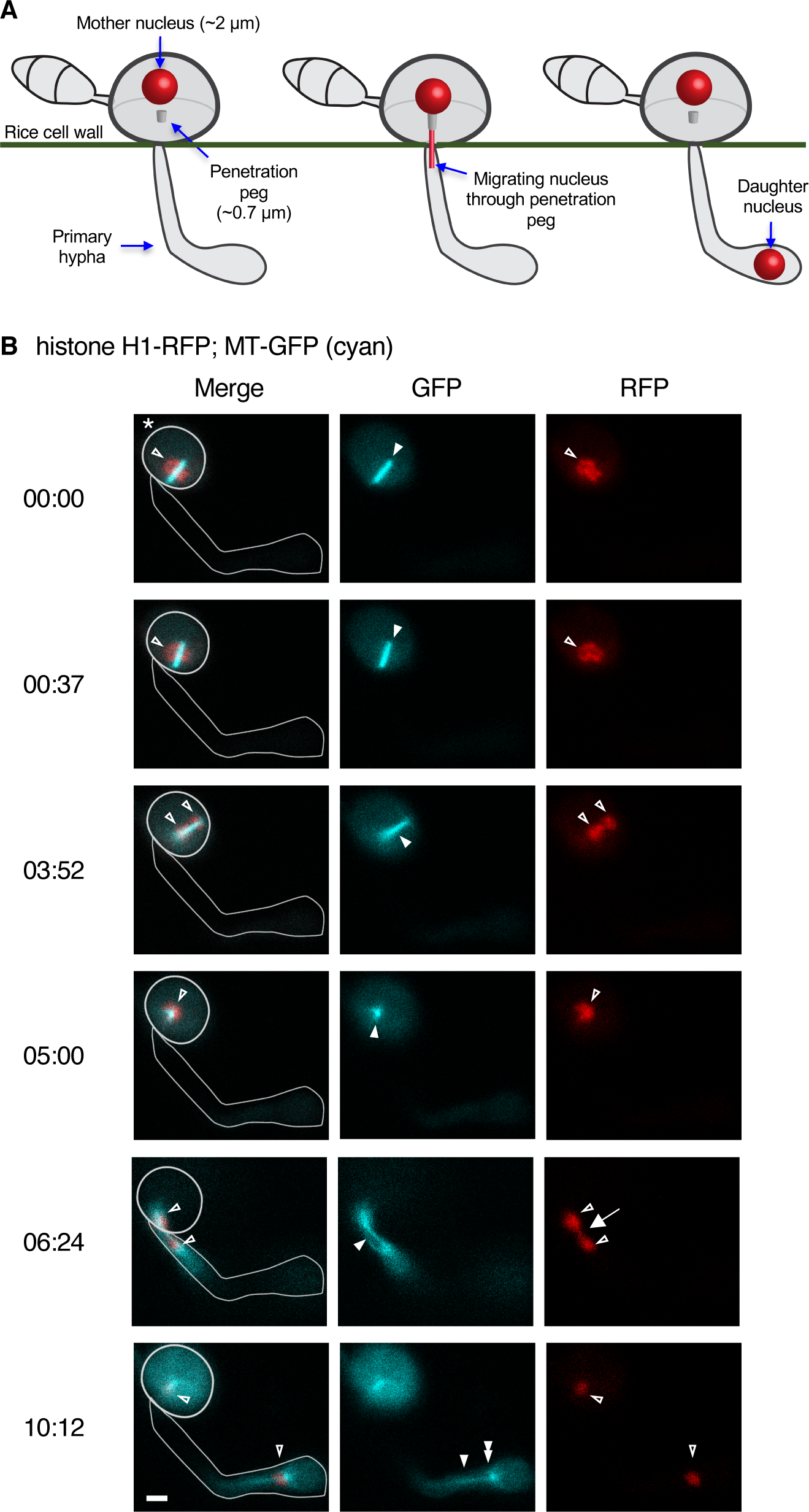
The spindle mediates nuclear migration through the penetration peg. (A) Schematic representation of extreme nuclear migration in *M. oryzae* during movement through the penetration peg (23, 24). The nucleus in the appressorium is referred to, here, as the mother nucleus (left). The appressorium forms on the surface of the rice leaf. The migrating nucleus, called the daughter nucleus, becomes elongated as it moves through the penetration peg (middle). Note that the penetration peg spans the rice cell wall, and does not protrude into the appressorium as shown in this depiction. The daughter nucleus is then positioned at the apical region of the primary hypha (right). The primary hypha forms inside the first-invaded rice epidermal cell. (B) Extreme nuclear migration through the penetration peg in *M. oryzae* strain CKF3578. The nucleus is shown in red (histone H1-RFP), and the spindle is shown in cyan (MT-GFP). Times is in minutes: seconds. An overlay to outline the appressorium and the primary hypha is provided in the merged channel. The GFP and RFP channel micrographs are purposively left without an overlay to more clearly display annotations. (00:00) The spindle (filled arrowhead) bisects the mother nucleus (open arrowhead) within the appressorium (asterisk). (00:37) The spindle rotates to become aligned for movement through the penetration peg. (03:52) Condensed chromosomes (chromatids) move towards the polar edges of the spindle (open arrowheads) while the spindle continues to become aligned for movement through the penetration peg. (05:00) the spindle (filled arrowhead) and mother nucleus (open arrowhead) are positioned for movement through the penetration peg. Due to the three-dimensional nature of the appressorium, the mother nucleus and spindle appear to be co-localized at this point. (06:24) The mother nucleus remains within the appressorium (top open arrowhead), while the daughter nucleus begins to separate from the mother nucleus (bottom open arrowhead) in the penetration peg, and is stretched within the penetration peg (arrow). The spindle (filled arrowhead) propels the daughter nucleus forward. (10:12) The mother nucleus is located within the appressorium (top open arrowhead), the daughter nucleus is delivered to the apical region of the primary hypha (bottom open arrowhead) by the spindle (filled arrowhead). The daughter bound spindle pole body is evident (double filled arrow). Micrographs are single informative focal planes. Scale bar is 2 µm.

**Fig. 2.**
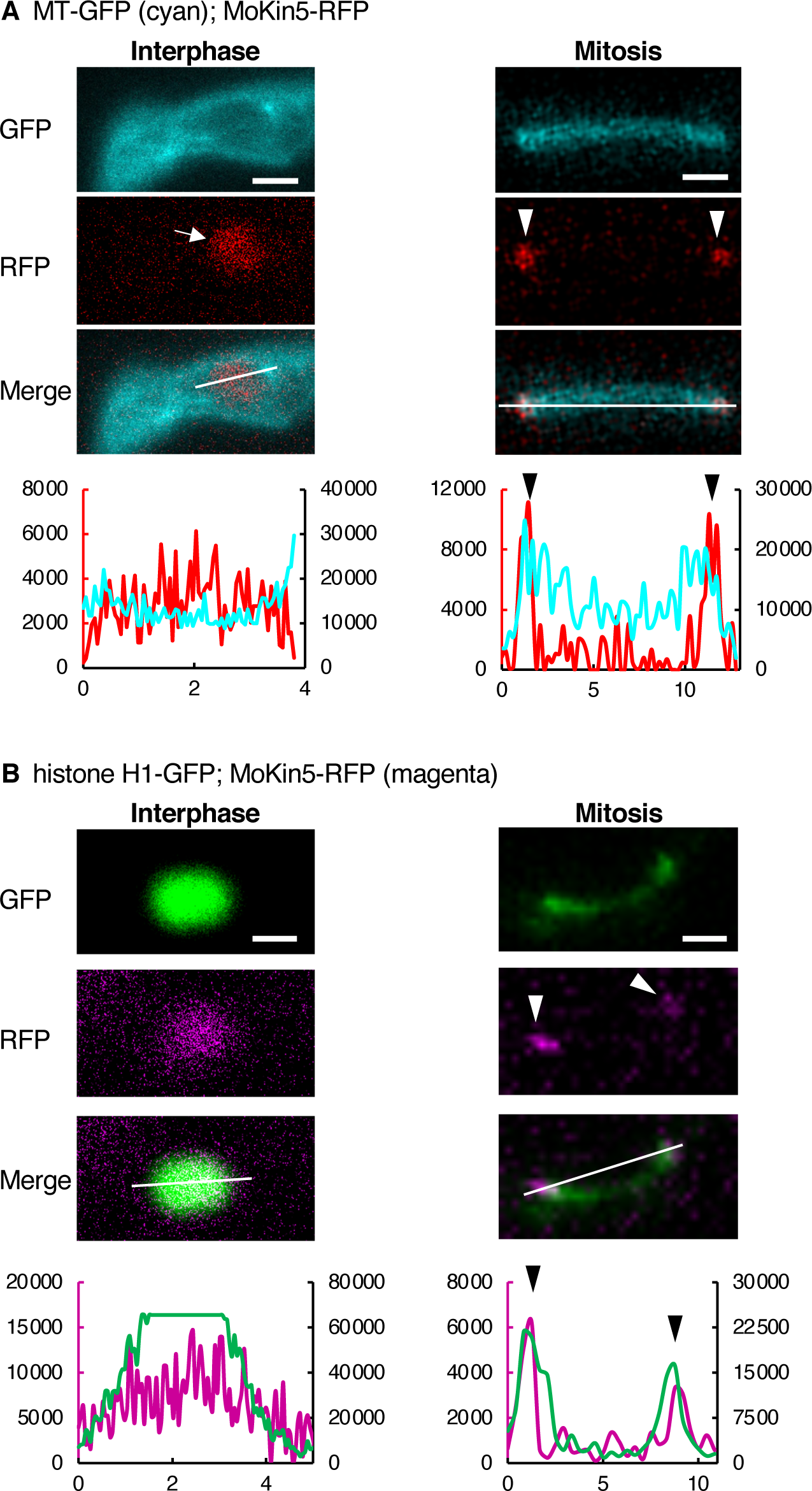
The localization patterns of MoKin5-RFP within invasive hyphae during interphase and mitosis in wild-type. All micrographs are single informative focal planes. Scale bars are 2 µm. Corresponding linescans quantify the fluorescence intensity in micrograph above. RFP intensity is displayed on primary vertical axis, GFP intensity is displayed on secondary vertical axis. Distance in µm is shown on horizontal axis. (A) The subcellular localization patterns of MoKin5-RFP relative to MT-GFP (cyan) in *M. oryzae* strain CKF4168. In interphase, MoKin5-RFP accumulates in the nucleus (left panel, arrow). During mitosis, MoKin5-RFP accumulates at the ends of the spindle (right panel, arrowheads). (B) The subcellular localization patterns of MoKin5-RFP relative to histone H1-GFP in *M. oryzae* strain CKF4208. In interphase, MoKin5-RFP co-localizes with histone H1-GFP (left panel). During mitosis, MoKin5-RFP accumulates at the ends of a dividing nucleus (right panel, arrowheads).

Despite evidence that the spindle is involved in nuclear migration at other *M. oryzae* infection stages, there is no direct evidence that the spindle mediates extreme nuclear migration through the penetration peg during initial rice cell colonization. Moreover, kinesin-5 and kinesin-14 motor proteins are yet to be discovered within *M. oryzae*. The goal of this study was twofold. First, we determined that the spindle is involved in nuclear migration through the penetration peg using confocal live-cell imaging of this remarkable cellular phenomenon. Second, we identified kinesin-5 and kinesin-14 motor proteins in *M. oryzae*. Identification of kinesin-5 and kinesin-14 in *M. oryzae* allowed us to develop an approach to genetically perturb spindle function specifically during extreme nuclear migration through the penetration peg. Our live-cell imaging observations coupled with experiments genetically perturbing spindle function demonstrate that the spindle mediates nuclear migration through the penetration peg.

## Results

### Dynamics of the spindle and the nucleus during extreme nuclear migration

We determined that the spindle formed and elongated during nuclear migration through the penetration peg using confocal live-cell imaging of a fluorescent *M. oryzae* strain. In this strain, microtubules (MTs) were labeled with β-tubulin-GFP (pseudocolored cyan throughout figures), and the nucleus was labeled with histone H1-tdTomato (RFP). We inoculated the fungal strain onto susceptible rice sheaths and observed nuclear migration through the penetration peg (Fig. 1B). Prior to nuclear migration through the penetration peg, the spindle bisected the mother nucleus within the appressorium (Fig. 1B, 00:00). The spindle and the mother nucleus rotated to become aligned to the axis of the appressorial pore and penetration peg (Fig. 1B, comparing 00:00 to 03:52). During migration through the penetration peg, the daughter nucleus stretched and separated (Fig. 1B, 06:24; *See Fig. 3A*). The mother nucleus remained within the appressorium. The daughter nucleus was delivered to the apical region of the primary hypha as the spindle elongated (Fig. 1B, comparing 06:24 to 10:12). During this nuclear migration event, the daughter nucleus traveled a total of 22µm from the site where the spindle first bisected the mother nucleus in the appressorium to the primary hypha. Consequently, nuclei migrating through the penetration peg undergo a longer nuclear migration compared to nuclei migrating in other IH cells. For instance, during nuclear migration in leading IH in wild-type, the maximum spindle length is typically less than 14 µm (*See Fig. 10E*). From these data, we concluded the spindle is involved in extreme nuclear migration through the penetration peg in wild-type.

**Fig. 3.**
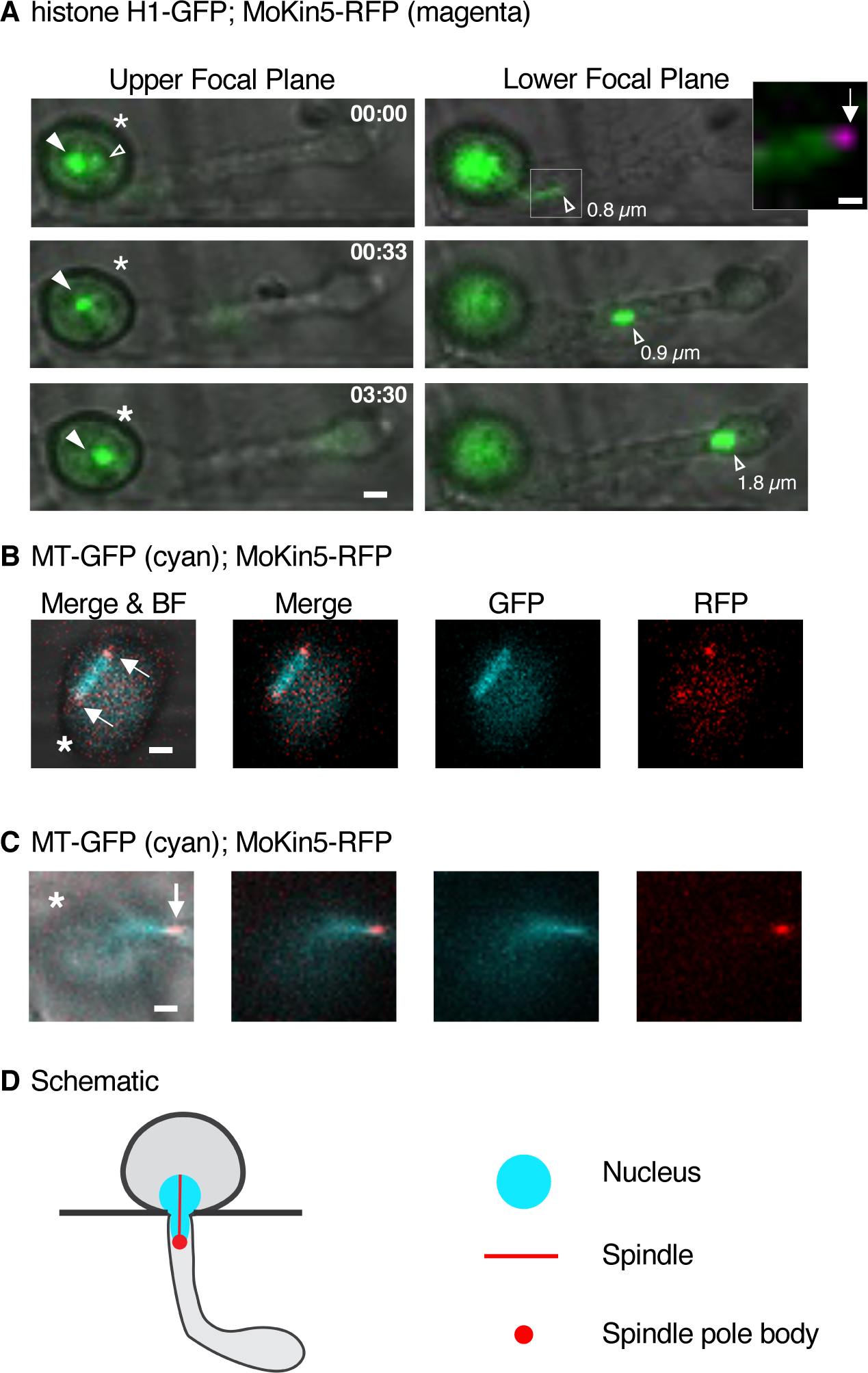
Nuclear, spindle, and spindle pole body dynamics during nuclear migration through the penetration peg in wild-type. All micrographs are single informative focal planes. All scale bars are 2 µm, except for the inset in panel 3A (far right; inset scale bar is 0.5 µm). Asterisks indicate the appressorium. Time is in minutes: seconds. (A) The dynamics of the mother and daughter nucleus (histone H1-GFP) during extreme nuclear migration through the penetration peg in *M. oryzae* strain CKF4208. Two informative focal planes are shown. Micrographs show GFP and brightfield channels. Upper focal plane panels (left) show the localization of the mother nucleus (filled arrowhead) within the appressorium. The mother nucleus remains within the appressorium throughout the event. (00:00) The migrating daughter nucleus is localized within the penetration peg (open arrowhead). The lower focal planes show the dynamics of the daughter nucleus (open arrowhead). As the daughter nucleus moves towards the apical region of the primary hypha, it expands in the y-dimension diameter. The inset (far right) corresponds to the white box in the lower focal plane image. The inset is a merged micrograph showing both the GFP and RFP channels. MoKin5-RFP (magenta) localizes at the tip of the dividing nucleus (arrow) as it migrates through the penetration peg. (B) MoKin5-RFP marks the spindle pole bodies (arrows) at the ends of the spindle (MT-GFP (cyan)) within the appressorium of *M. oryzae* strain CKF4168. (C) The daughter bound spindle pole body (marked by MoKin5-RFP; arrow) localizes at the tip of the spindle (MT-GFP) during movement through the penetration peg in *M. oryzae* strain CKF4168. (D) Schematic representation of the position of the nucleus, spindle, and daughter bound spindle pole body during movement through the penetration peg in wild-type.

### Identification of MoKin5-RFP as a maker for spindle pole bodies (SPBs) during mitosis

Since kinesin-5 plays an important role in formation and maintenance of spindles in other fungi, we identified the kinesin-5 homolog, *MoKin5*, in *M. oryzae*. *MoKin5* (MGG_01175) was identified based on protein sequence homology to previously characterized kinesin-5 proteins in *Aspergillus nidulans, Schizosaccharomyces pombe,* and *Saccharomyces cerevisiae* (Fig. S1). MoKin5 shared 62% global similarity to BimC (AN3363), which is kinesin-5 in *A. nidulans*. Importantly, MoKin5 contained a predicted kinesin motor domain near its N-terminus. The location of this kinesin motor domain is characteristic of kinesin-5 motor proteins, and indicate that MoKin5 likely walks towards the plus-ends of MTs (9). We cloned MoKin5 to produce a *MoKin5*-tdTomato (RFP) construct driven by the native *MoKin5* gene promoter. We generated *M. oryzae* fluorescent strains to determine the subcellular localization of MoKin5-RFP in wild-type during interphase and mitosis.

The subcellular localization of MoKin5-RFP was first determined relative to MT-GFP during interphase in IH using live-cell confocal imaging. During interphase, MTs were arranged in a cage-like manner around a mass of MoKin5-RFP fluorescence (Fig. 2A). We hypothesized that the mass of MoKin5-RFP fluorescence represented the nucleus. We confirmed that MoKin5-RFP localized within the nucleus during interphase in an additional *M. oryzae* strain. This strain expressed MoKin5-RFP and histone H1-GFP to label the nucleus. During interphase, MoKin5-RFP and histone H1-GFP co-localized (Fig. 2B). We concluded that when MoKin5-RFP is expressed from its native promoter it accumulates within the nucleus during interphase.

During mitosis, the localization of MoKin5-RFP changed in wild-type strains. MoKin5-RFP accumulated at the ends of the spindle (MT-GFP) (Fig. 2A). We observed dividing nuclei (histone H1-GFP) and found MoKin5-RFP accumulated at the polar ends of the mitotic nucleus (Fig. 2B). These data suggested MoKin5-RFP localized at the spindle pole bodies (SPBs) during mitosis (Fig. 2). We corroborated this finding by comparing the subcellular localization of MoKin5-RFP to a known component of SPBs, γ-tubulin. We identified *M. oryzae γ-tubulin* (MGG_00961) based on protein homology to *A. nidulans γ-tubulin* (AN0676, MipA; Fig. S2). We generated a reporter strain expressing γ-tubulin-RFP and MT-GFP. Comparing the localization of MoKin5-RFP to the localization of γ-tubulin-RFP relative to the spindle during appressorium development revealed identical subcellular localization patterns at the ends of the spindle (Fig. S2). Taken together, these data showed that MoKin5-RFP accumulated at the SPBs during mitosis. MoKin5-RFP was subsequently utilized as a reporter for the SPBs during mitosis.

### The dynamics of spindle pole bodies (SPBs) relative to the nuclei and spindle during extreme nuclear migration

We investigated the arrangement of the mother and daughter nuclei in relation to the SPBs during extreme nuclear migration through the penetration peg using live-cell confocal microscopy of a fluorescent fungal strain infecting rice. In this *M. oryzae* strain, the nucleus was labeled with histone H1-GFP and SPBs were labeled with MoKin5-RFP. We observed that the mother nucleus remained within the appressorium, while the migrating daughter nucleus was delivered to the apical region of the primary hypha (Fig. 3A). The daughter nucleus became highly elongated as it migrated through the penetration peg (Fig. 3A, 00:00, Lower Focal Plane). MoKin5-RFP, marking the SPB bound to the migrating daughter nucleus, was localized at the apical tip of the elongated daughter nucleus during movement through the penetration peg (Fig. 3A, 00:00, Lower Focal Plane, Inset). The SPB bound to the mother nucleus was not detectable in our microscopy, possibly due to relatively strong autofluorescence in the melanized appressorium. The y-dimension diameter of the daughter nucleus expanded throughout the nuclear migration event. The diameter of the apical tip of the daughter nucleus was ∼0.8 µm immediately following movement through the penetration peg (Fig. 3A,00:00, Lower Focal Plane) but increased to ∼1.8 µm as it neared the apical region of the primary hypha (Fig. 3A; 03:30, Lower Focal Plane). We also followed the dynamics of the SPBs in relation to the spindle during extreme nuclear migration through the penetration peg in an additional fluorescent *M. oryzae* strain. The spindle (MT-GFP) and SPBs (MoKin5-RFP) first formed within the appressorium (Fig. 3B). As expected, the daughter bound SPB proceeded the spindle during movement through the penetration peg (Fig. 3C). These data established the typical wild-type dynamics of the nucleus, spindle, and SPBs during extreme nuclear migration through the penetration peg (Fig. 3D; *See Fig. 11*).

### Development of an inducible promoter system to perturb spindle function during nuclear migration through the penetration peg

Our observations in wild-type pointed to the importance of the spindle in mediating extreme nuclear migration through the penetration peg. We hypothesized that genetically perturbing spindle function would impair nuclear migration at this infection stage. Yet we lacked an inducible promoter system to test this hypothesis. To overcome this limitation, we exploited the effector biology of *M. oryzae*. Effectors are small proteins secreted by pathogens to modulate their hosts during infection (27). We developed an inducible promoter system using the promoter of the *M. oryzae* effector gene, *Bas4*. Bas4 is an apoplastic effector whose promoter activity is highly induced upon initial penetration into plant tissue (28, 29). We reasoned that we could generate an inducible overexpression construct by expressing a target gene with the *Bas4* promoter (*p*). The first inducible overexpression construct we generated contained the *Bas4p* fused to the *MoKin5* coding sequence and accompanying terminator region (*Bas4p-MoKin5*; Fig. 4A).

**Fig. 4.**
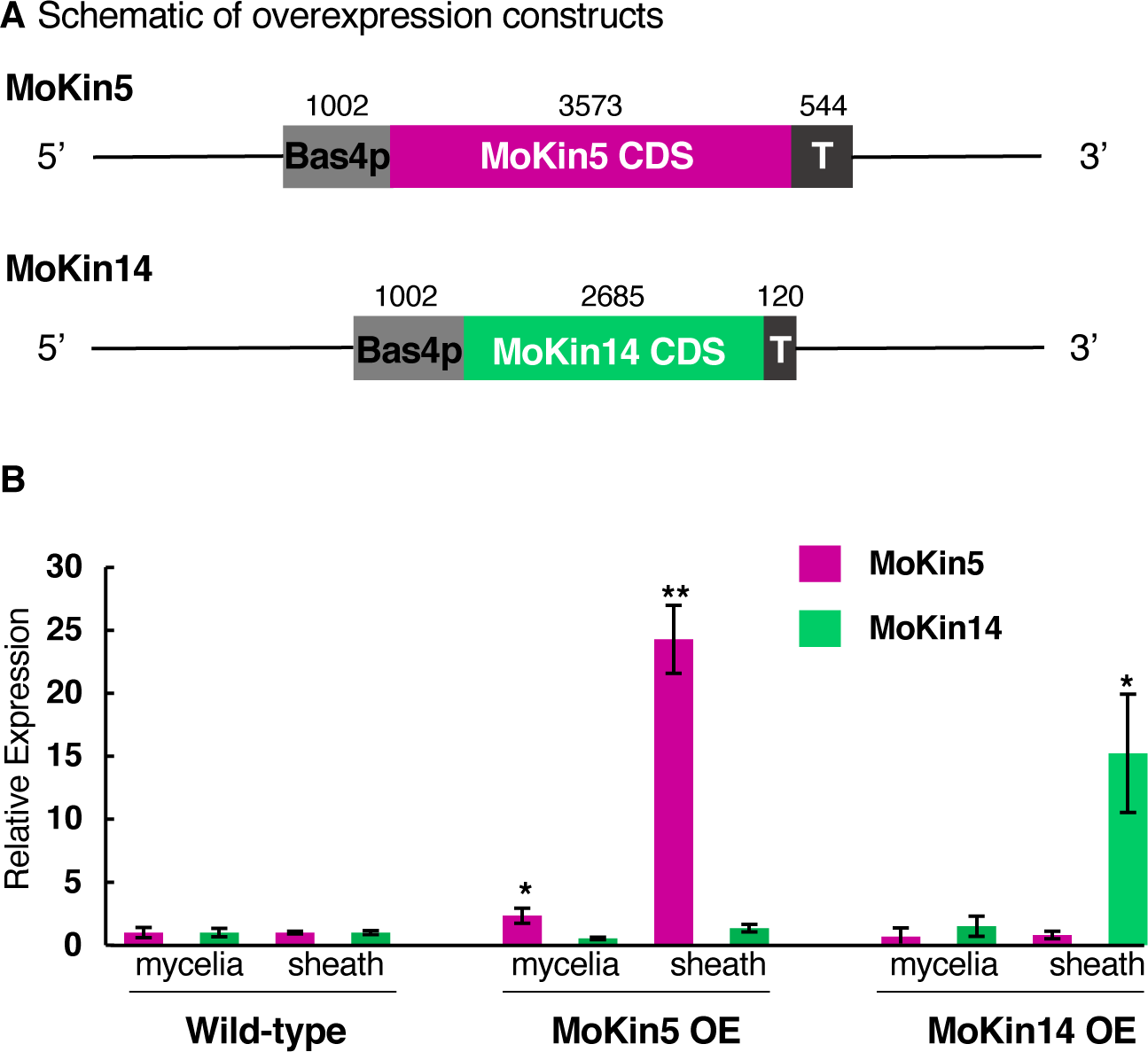
Relative expression levels of *MoKin5* and *MoKin14* driven by the *Bas4* promoter. (A) Schematics of *MoKin5* and *MoKin14* overexpression (OE) constructs. (B) Relative expression of *MoKin5* (magenta) and *MoKin14* (green) in wild-type (CKF3578), MoKin5 OE (CKF4108), and MoKin14 OE (CKF4106) mycelia and YT16-infected rice sheaths. Data from two separate RT-qPCR experiments are shown. Samples were normalized relative to *actin* in each strain. Mycelia were harvested after 5 days growth in complete medium. Infected YT16 rice sheaths were harvested at 30-31 hours post inoculation. Significance was determined using a Student’s t-test assuming unequal variance. P-value of *MoKin5* in MoKin5 OE mycelia is 0.04. P-value of *MoKin5* in MoKin5 OE sheaths is 0.003. P-value of *MoKin14* in MoKin14 OE sheath is 0.03. Error bars are 95% confidence intervals.

We conducted RT-qPCRs to determine the expression of *MoKin5* relative to *actin* in two fungal cell types: vegetative mycelia and within IH growing inside the first-invaded rice cell. In wild-type, the *MoKin5* expression level relative to *actin* was 1 in both mycelia (± 0.4 margin of error) and IH (± 0.1 margin of error) (Fig. 4B). In the fungal strain carrying the *Bas4p-MoKin5* construct, *MoKin5* expression relative to *actin* was 2.3 (± 0.6 margin of error) in mycelia. In the rice sheath samples infected by the same fungal strain, the relative expression of *MoKin5* to *actin* was 24.3 (± 2.7 margin of error). The relatively high expression of *MoKin5* within IH validated the use of the *Bas4* promoter to induce overexpression of a gene during early rice infection. *Magnaporthe oryzae* strains carrying a *Bas4p-MoKin5* construct were therefore referred to as MoKin5 overexpression (OE) strains.

### MoKin5 OE causes defects in nuclear morphology and positioning

The development of the *Bas4p* inducible overexpression system allowed us to test our hypothesis that genetically perturbing spindle function would impair nuclear migration through the penetration peg. We reasoned that one consequence of impaired spindle function would be disruption in nuclear positioning within the appressorium and primary hypha relative to wild-type. We conducted live-cell confocal microscopy of fluorescent fungal strains expressing histone H1-RFP to label nuclei infecting a susceptible rice cultivar at two timepoints, ∼28 hpi (early) and ∼48 hpi (late). At the early timepoint, a majority of infection sites displayed a single nucleus within the appressorium and a single nucleus within the primary hypha in wild-type (Fig. 5A; Fig. S3). Nuclear positioning within the MoKin5 OE strain was highly disrupted. In the MoKin5 OE strain, only 2% (n=2) of infection sites displayed a single nucleus within the appressorium and a single nucleus within the primary hypha at the early time point (Fig. 5A). In the MoKin5 OE strain, 25% (n=31) of infection sites displayed an anucleate appressorium with a single enlarged nucleus within the primary hypha at the early timepoint (Fig. 5A, Fig. S3A). This phenotype was especially striking because all the infection sites scored contained intact appressoria. That is, any infection site that showed a collapsed appressorium in the bright-field channel was excluded from analysis. Additional defects in nuclear morphology and positioning were observed in the MoKin5 OE strains at the early and late timepoints (Fig. S3). Prominent defects included nuclear fragments within the appressorium (Fig. 6B), nuclear fragments within the appressorium and primary hypha (Fig. 6C), nuclear fragments exclusively within the primary hypha (Fig. 7), and a single enlarged nucleus that appeared to be stuck within the penetration peg (Fig. S3C). We concluded that MoKin5 OE caused failure in extreme nuclear migration through the penetration peg.

**Fig. 5.**
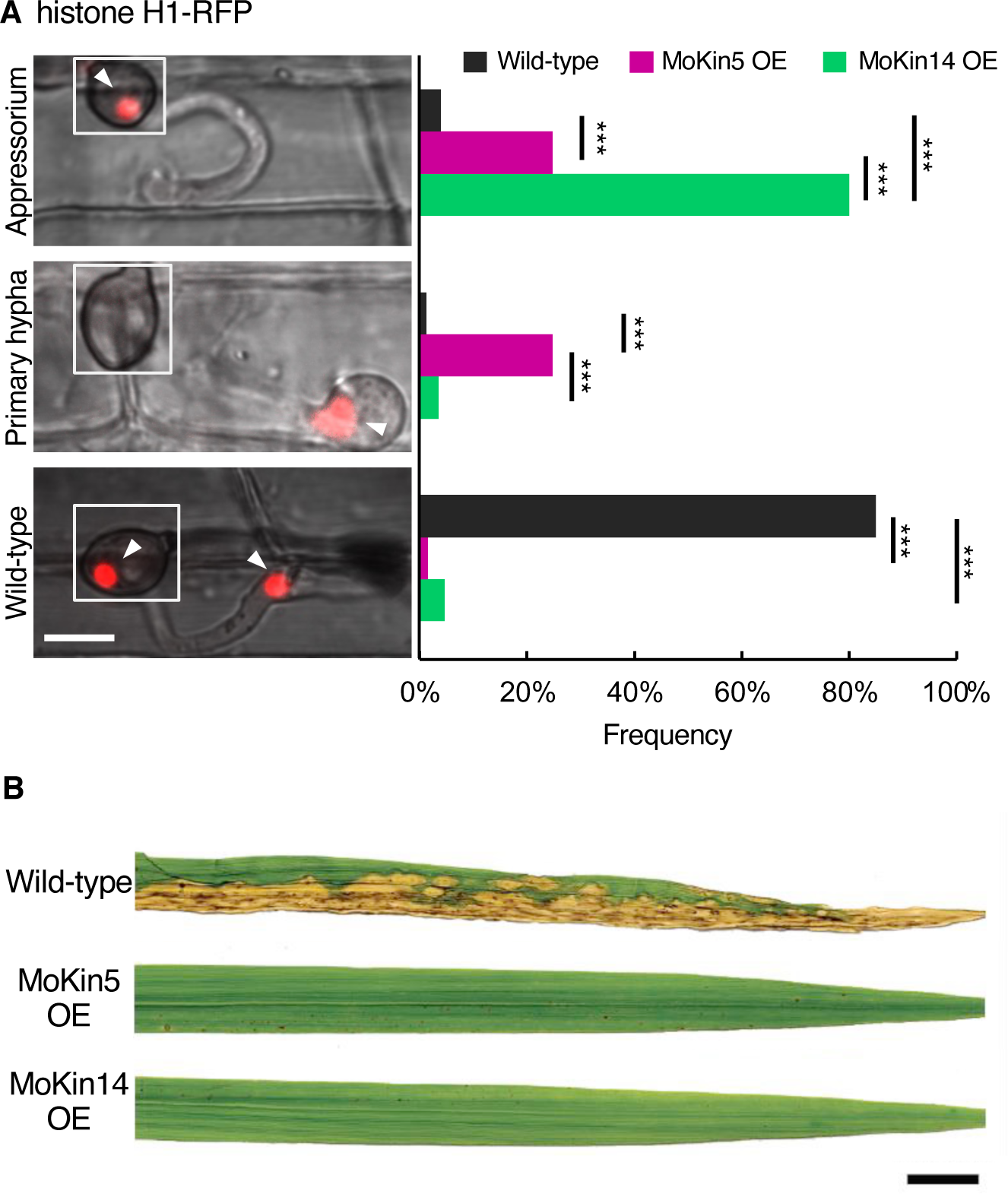
MoKin5 OE and MoKin14 OE cause defects in nuclear positioning and morphology, and decreases in virulence on rice. (A) Frequency of most commonly observed nuclear positioning and morphology phenotypes at ∼28 hours post inoculation in wild-type (CKF3578 and CKF3971, n=153), MoKin5 OE (CKF4108, n=125), and MoKin14 OE (CKF4106 and CKF4093, n=85) strains. Example micrographs on the left are single focal planes. Only histone-H1 and brightfield channels are shown in example micrographs. The inset shows a single focal plane depicting the position of the mother nucleus within the appressorium. Scale bar is 5 µm. Top panel shows a single nucleus within the appressorium (arrowhead). The middle panel shows a single nucleus within the primary hypha (arrowhead). The bottom panel shows a single nucleus within the appressorium (arrowhead) and a single nucleus within the primary hypha (arrowhead). Statistical significance of nuclear phenotype frequency was determined using two-tailed Fisher’s exact tests. *** represent p-values less than 0.0001. Non-significant p-values are 0.22 for MoKin5 OE relative to MoKin14 OE in the wild-type category and 0.35 for wild-type relative to MoKin14 OE in the primary hypha category. The frequency of all observed nuclear phenotypes at the early timepoint is available in Fig. S3A. (B) Images of representative infected leaves from whole-plant spray inoculations. Wild-type is CKF3578, MoKin5 OE is CKF4108, and MoKin14 OE is CKF4106. Scale bar is 1 cm.

**Fig. 6.**
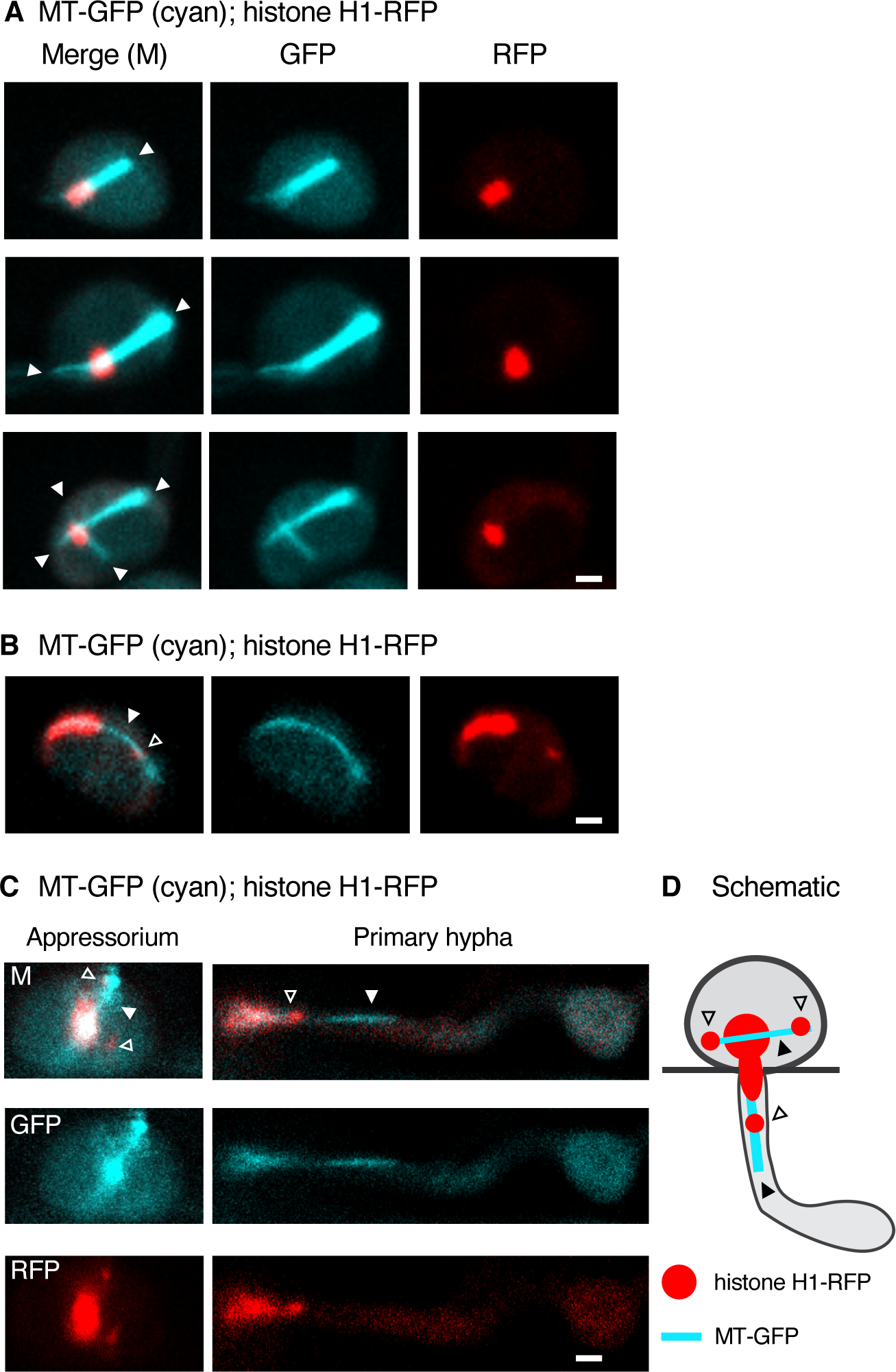
Formation of microtubule (MT) protrusions and nuclear fragments within the MoKin5 OE strain CKF4108. All micrographs are informative single focal planes, except Fig. 6C (appressorium, left panels are maximum intensity projections). All scale bars are 2 µm. (A) Single (top), double (middle), and triple + (bottom) MT protrusions (filled arrowhead) within appressoria. (B) A nuclear fragment (open arrowhead) on a MT protrusion (filled arrowhead) within an appressorium. (C) The arrangement of the nucleus, nuclear fragments, and MT protrusions during extreme nuclear migration through the penetration peg. Within the appressorium (left panels) two nuclear fragments (open arrowheads) form along MT protrusions. A single MT protrusion is in focus (filled arrowhead), while another MT protrusion is out of focus (not marked). The micrograph on the right shows a lower single informative focal plane. Here, an MT protrusion (filled arrowhead) leads the nuclear fragment (open arrowhead) just emerging from the penetration peg. (D) A two-dimensional schematic representation of the dynamics presented in Fig. 6C.

**Fig. 7.**
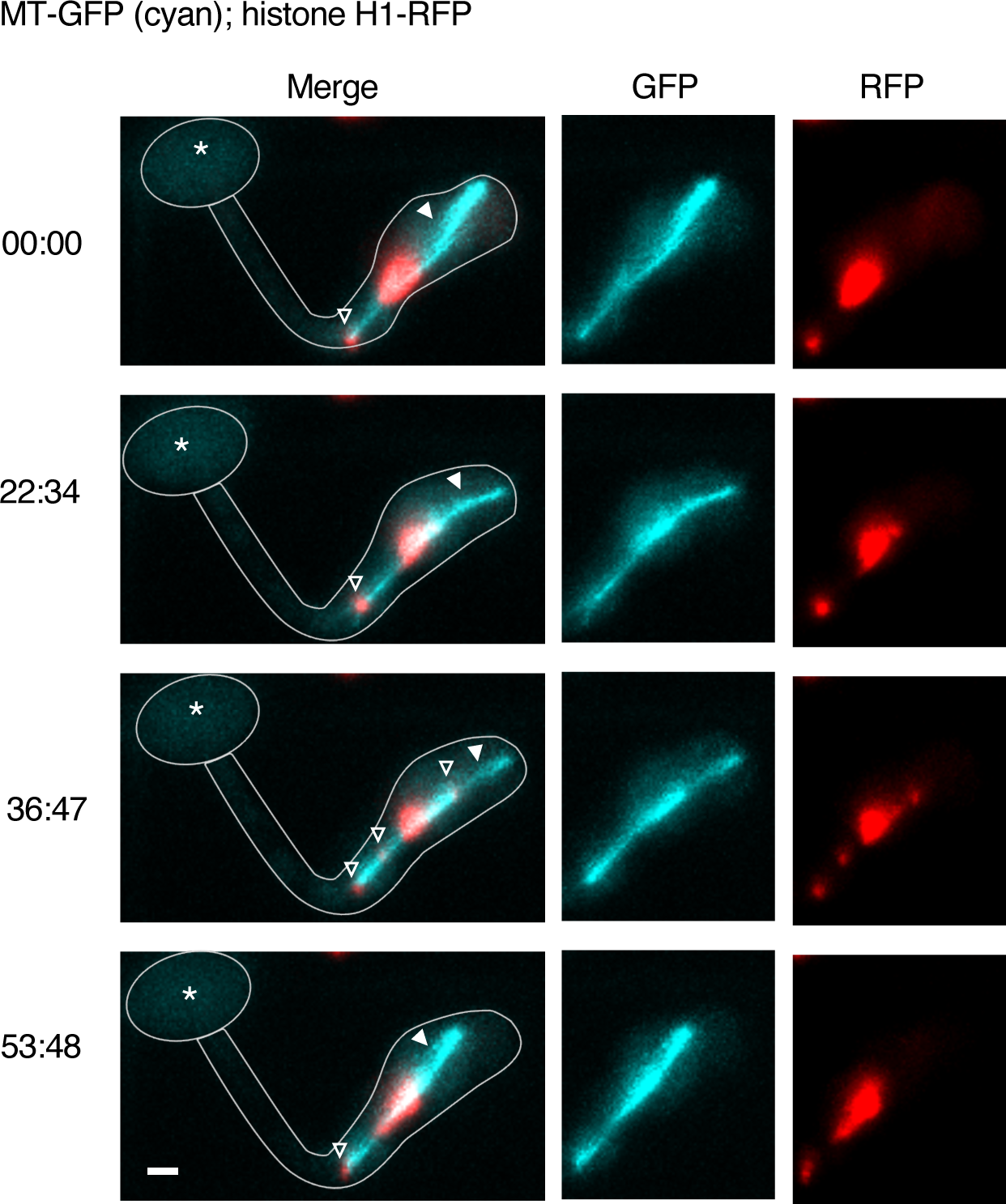
The behavior of small nuclear fragments (histone H1-RFP, open arrowheads) along the spindle and MT protrusion (MT-EGFP (cyan); filled arrowhead) in a primary hypha of the MoKin5 OE strain CKF4108. Each micrograph is a maximum intensity projection of informative single focal planes. The scale bar is 2 µm. Asterisks indicate an anucleate appressorium. An overlay outlining the appressorium and primary hypha is present in the merged channel micrographs. Time is in minutes: seconds.

### MoKin5 OE causes defects in fungal development and virulence on rice

Considering the dramatic defects in nuclear morphology and positioning in the MoKin5 OE strain, we determined the effect of MoKin5 OE on IH development and blast lesion development on whole rice plants. At the late timepoint (∼48 hpi), MoKin5 OE strains typically failed to develop beyond the primary hyphal stage of development (Fig. S4). We conducted whole-plant spray inoculations to determine if MoKin5 OE strains retained virulence of rice. In whole-plant spray inoculations, the mean percentage of diseased tissue area was 68% (±13% margin of error) in wild-type. The mean percentage of diseased tissue was 0% in the MoKin5 OE strain (representative infected leaves in Fig. 5B). We concluded that MoKin5 OE strains failed to develop beyond the primary hyphal stage within the first-invaded rice cell, which caused a drastic reduction in virulence on rice.

### MoKin5 OE leads to the formation of MT protrusions and nuclear fragments

The severe developmental defects caused by MoKin5 OE warranted a mechanistic explanation. We investigated the effect of MoKin5 OE upon the spindle during nuclear migration through the penetration peg using live-cell confocal imaging of a MoKin5 OE strain infecting rice sheaths. We initiated our investigation by examining the dynamics of the spindle (MT-GFP) relative to the nucleus (histone H1-RFP) in the appressoria of a MoKin5 OE strain at the start of mitosis. In these MoKin5 OE infection sites, a bar of MT-GFP extended in an abnormal and persistent manner beyond the circumference of the nucleus, as recognized by histone H1-RFP fluorescence, in appressoria and primary hyphae (Fig. 6A). These persistent MT-GFP structures, which we refer to as MT protrusions, were not observed in wild-type. We quantified the frequency of single, double, and triple+ MT protrusions relative to the nucleus within appressoria (Fig. 6A, n=22). We found that 45% (n=10) of infection sites contained a single MT protrusion (Fig. 6A, top panel), 36% (n=8) of infection sites contained double MT protrusions (Fig. 6A, middle panel), and 18% (n=4) of infection sites contained three or more MT protrusions (Fig. 6A, bottom panel). We concluded that MoKin5 OE did impair spindle function by preventing formation of a typical spindle within the appressorium.

Additional MT protrusion and nuclear positioning phenotypes were observed in the MoKin5 OE strain within appressoria and primary hyphae. In appressoria, small nuclear fragments were distributed along the MT protrusions (Fig. 6B-C). We followed the relative position of the spindle/MT protrusions, and the nucleus/nuclear fragments during extreme nuclear migration through the penetration peg in the MoKin5 OE strain. In the MoKin5 OE strain, the MT protrusion proceeded the nuclear fragment (Fig. 6C-6D). This arrangement was in stark contrast to the arrangement of the spindle and nucleus in wild-type (Fig. 3C-3D). Nuclear fragments tended to occur more frequently within the primary hyphae of the MoKin5 OE strain. At the early timepoint, only 7% (n=9) of MoKin5 OE infection sites showed nuclear fragments in the appressorium, whereas 12% (n=15) of MoKin5 OE infection sites showed nuclear fragments in the primary hypha exclusively (Group 2 vs Group 4 in Fig. S3A). We conducted time-lapse confocal microscopy to further investigate the nature of the MT protrusions and nuclear fragments within the primary hyphae. Within primary hyphae, nuclear fragments separated and merged over time along the MT protrusion (Fig. 7). From these data, we concluded that MoKin5 OE caused formation of MT protrusions. These MT protrusions contributed to the formation of nuclear fragments beginning within the appressorium, and that the nuclear fragments merged together to form a single enlarged nucleus within the primary hypha.

### MoKin5 OE causes defects in spindle polarity

We pursued a mechanistic understanding of how the MT protrusions observed in the MoKin5 OE strain contributed to formation of nuclear fragments within the appressorium. We conducted confocal microscopy of appressoria in an *M. oryzae* strain co-expressing three constructs: *MoKin5*-RFP driven from its native promoter, *Bas4p-MoKin5*, and *β-tubulin*-GFP to label the spindle. During mitosis in the appressoria of the MoKin5 OE strain, MoKin5-RFP localized along the spindle (Fig 8A; Fig S5). The MoKin5-RFP localization pattern in the MoKin5 OE strain differed dramatically from wild-type during mitosis (Fig. 2; Fig. 3B). In wild-type, MoKin5-RFP accumulated only at the SPBs. We concluded that during mitosis MoKin5 OE caused MoKin5-RFP to inappropriately localize along MTs within the spindle.

**Fig. 8.**
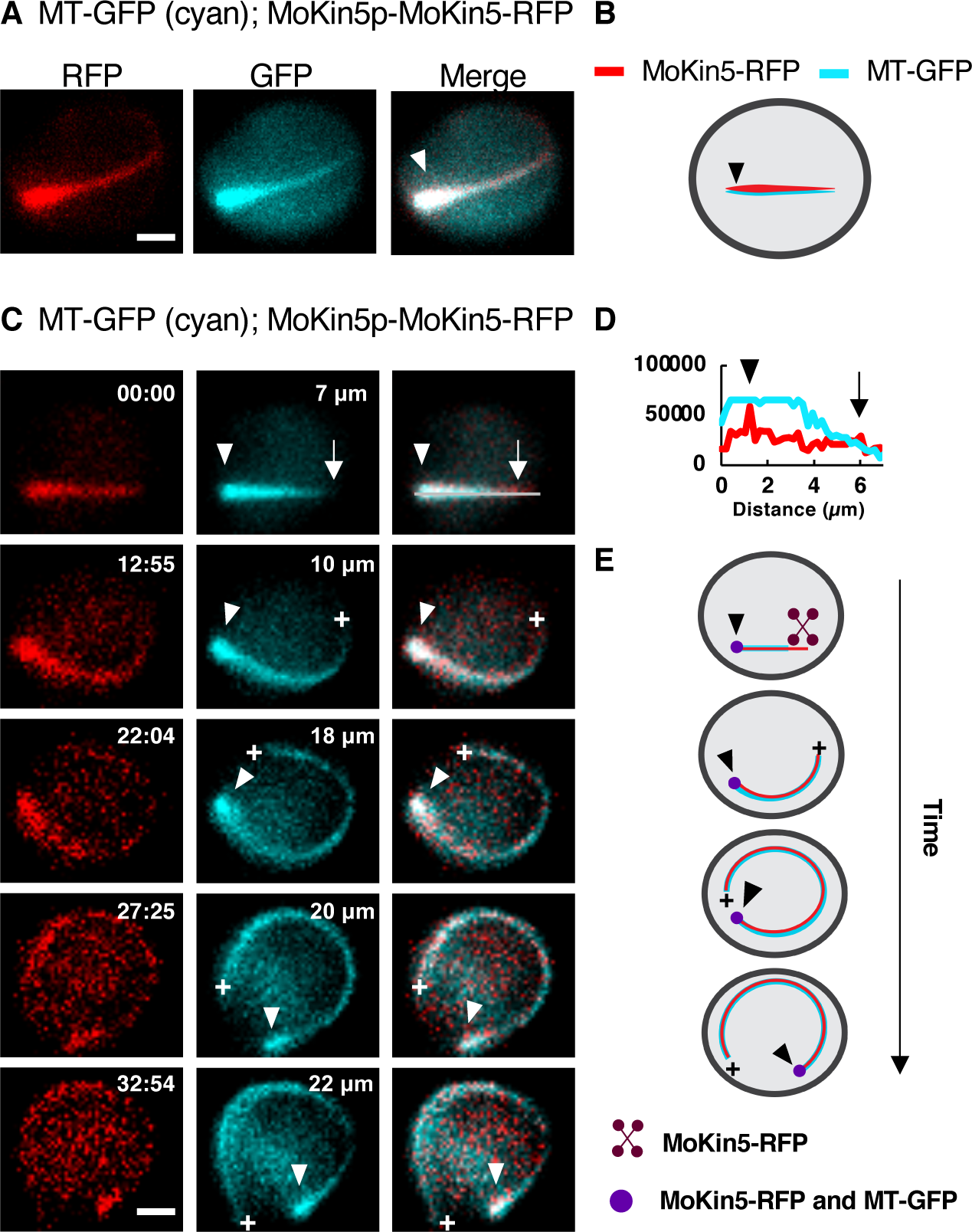
The localization of MoKin5-RFP relative to MT-GFP (cyan) in spindles located within the appressoria of MoKin5 OE strain CKF4203. Maximum intensity projections of informative single focal planes are shown. Scale bars are 2 µm. (A) A bright focus (filled arrowhead) of MT-GFP and MoKin5-RFP is found at one end of the MoKin5 OE spindle. (B) A schematic representation of the patterns shown in Fig. 8A. (C) Time-lapse micrograph series showing spindle elongation in the MoKin5 OE strain. Time is in minutes: seconds. (00:00) The bright focus of MT-GFP (cyan) and MoKin5-RFP (filled arrowhead) is opposite a small and transient accumulation of MoKin5-RFP at the other end of the spindle (arrow). The initial length of the MoKin5 OE spindle is 7 µm. Over time, the spindle grows from a single end (plus symbol). The MoKin5 OE spindle elongates in a manner that follows the curvature of the appressorium, and the spindle rotates (compare 12:55 to 32:54). (D) Linescan quantifying fluorescence intensity of MT-GFP and MoKin5-RFP corresponding to timepoint 00:00 in Fig. 8C. MT-GFP and MoKin5-RFP signal shows an accumulation at one end of the spindle (black arrowhead). MoKin5-RFP shows a smaller accumulation at the opposite end of the spindle (black arrow). (E) Schematic representation of the MoKin5 OE spindle elongating within the appressorium based on data presented in Fig. 8C. The focus of MoKin5-RFP and MT-GFP at the end of the spindle is shown by a purple circle, and corresponds to the white arrowheads in Fig. 8C. MoKin5-RFP shows a brief accumulation at the opposite end of the spindle at the start of spindle elongation (top image). The spindle grows from the end marked by the plus symbol.

We continued to investigate the nature of the spindle and MoKin5-RFP in the MoKin5 OE strain by conducting time-lapse confocal microscopy of mitotic appressoria (Fig. 8C). In these appressoria we made several important observations. First, we observed that the MT-GFP and MoKin5-RFP signal displayed a relatively bright focus at one end of the spindle (Fig. 8, filled arrowheads in the merge channel). Second, the spindle elongated from only a single end, which we call the growing plus-end (Fig. 8C, Fig. 8E, plus symbol). This spindle elongation followed the curvature of the appressorium. Third, at the very early stage of spindle elongation, MoKin5-RFP showed a brief accumulation at the growing plus-end (Fig. 8C, 00:00, arrow; Fig. 8D, arrow). Finally, we observed that the MoKin5 OE spindle continued to elongate and rotate within the appressorium for at least 32 minutes (Fig. 8C). These data suggested that the MoKin5 OE spindle displays aberrant polarity likely due to a combination of excessive outward forces acting on the spindle and excessive polymerization of MTs within the spindle. Recently, monomeric human kinesin-5 was found to act as a promoter of MT polymerization at the plus-ends of MTs (30).

### MoKin14 OE causes defects in fungal development and virulence on rice

Due to the prominent defects in nuclear migration caused by MoKin5 OE, we investigated the effect of MoKin14 OE on extreme nuclear migration through the penetration peg. We identified a kinesin-14 motor protein in *M. oryzae* (*MoKin14*, MGG_05350) through protein homology to other known kinesin-14 proteins (Fig. S6). MoKin14 shared 55.9% similarity to KlpA (AN6340), which is kinesin-14 in *A. nidulans.* MoKin14 also contained a predicted kinesin motor domain at the C-terminus. This C-terminal kinesin motor domain is a characteristic of the kinesin-14 superfamily, and indicates that MoKin14 likely walks towards the minus-end of MTs (10). Like the MoKin5 OE strain, we generated several *M. oryzae* strains of MoKin14 OE (*MoKin14* under control of the *Bas4* promoter; *Bas4*p-*MoKin14*) for subsequent analysis of nuclear positioning, IH development, and virulence on rice.

Our first step in the analysis of the MoKin14 OE strains was to confirm that expressing *MoKin14* from the *Bas4* promoter caused an increase in the relative expression of *MoKin14* within IH (Fig. 4A). We conducted RT-qPCRs. In both mycelia and IH of wild-type, the expression of *MoKin14* relative to *actin* was 1 (± 0.3 in margin of error in mycelia and ± 0.2 margin of error in IH). In the *M. oryzae* strain carrying the *Bas4—MoKin14* construct, the expression of *MoKin14* relative to *actin* was 1.5 (± 0.6 margin of error) in mycelia and 15.2 (± 4.7 margin of error) in IH (Fig. 4B). Because we validated that a strain carrying the *Bas4p-MoKin14* construct did, indeed, cause an overexpression of MoKin14 in the early stages of rice infection, we then determined the effect of MoKin14 overexpression (OE) on nuclear positioning in appressoria and primary hyphae. At the early timepoint (∼28 hpi), 80% (n=68) of the MoKin14 OE sites displayed a single nucleus (histone H1-RFP) within the appressorium, which differed from both the wild-type and MoKin5 OE phenotypes (Fig. 5A; Fig. S3). The MoKin14 OE strains also showed a drastic arrest in IH development at ∼48 hpi. At this timepoint, MoKin14 OE strains were typically arrested at the primary hyphal stage of development (Fig. S4). We conducted whole-plant spray inoculations and found that the MoKin14 OE strain did not display virulence on rice. The mean percentage of diseased tissue area was 0% in the MoKin14 OE strain compared to 68% (±13% margin of error) in wild-type (representative leaves in Fig. 5B). From these results, we concluded that MoKin14 OE caused a failure in extreme nuclear migration through the penetration peg. The failure in extreme nuclear migration led to defects in IH development, which prevented virulence on rice.

### MoKin14 OE causes formation of monopolar spindles

MoKin14 OE caused failure in extreme nuclear migration through the penetration peg, and resulted in a nuclear positioning phenotype that was unique relative to wild-type and MoKin5 OE strains. This observation suggested that MoKin14 OE induced a distinct effect upon the spindle. We hypothesized that if MoKin14 generated an inward force on SPBs within *M. oryzae,* overexpressing MoKin14 with the *Bas4* promoter would cause formation of monopolar spindles within the appressorium. We conducted live-cell confocal microscopy of *M. oryzae* strains expressing MT-GFP and histone H1-RFP to determine the dynamics of the spindle in relation to the nucleus within the appressorium. Within appressoria, the spindle phenotype of the MoKin14 OE strain differed from both wild-type and the MoKin5 OE strain (Fig. 9). In wild-type, a spindle bisected the nucleus (Fig. 9A, top panel). We also detected the asynchronous movement of chromatids towards the ends of the spindle in wild-type (Fig. 9A, bottom panel). In contrast, MoKin14 OE resulted in a single focus of MT-GFP overlapping with the nucleus within the appressorium (Fig. 9B, top panel). MTs emanated from this single focus and, at times, the mother nucleus appeared to be arrested in mitosis. For example, within the MoKin14 OE strain, a “butterfly” shaped nucleus was observed within the appressorium, in which chromatids appear to be arrested in the process of dividing (Fig. 9B, bottom panel). Consistent with our previous observations, the MoKin5 OE spindle did not form a typical spindle, but instead appeared as a half spindle relative to the nucleus within the appressorium (Fig. 9C). We concluded that MoKin14 OE caused formation of monopolar spindles within the appressorium.

**Fig. 9.**
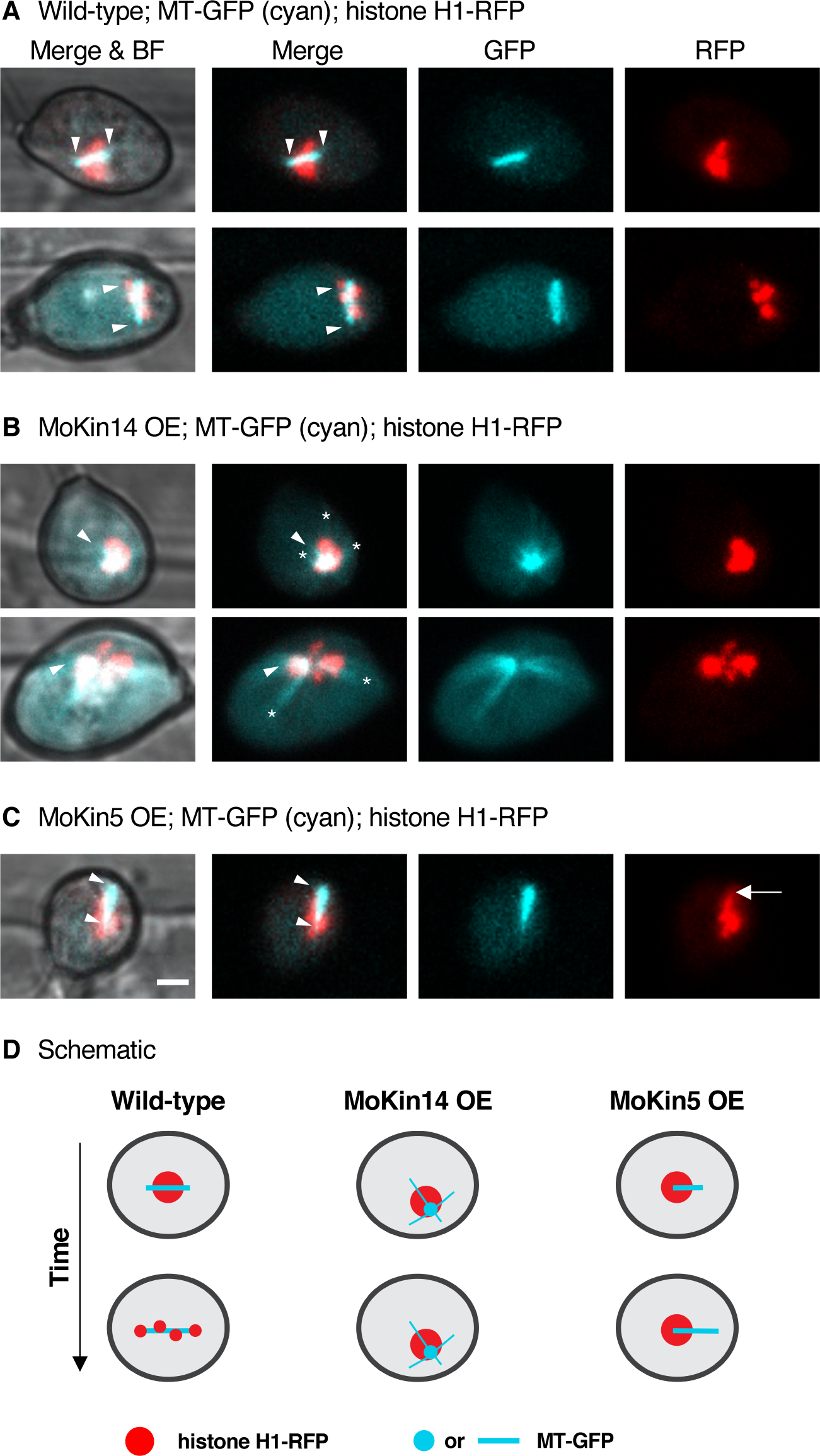
The arrangement of the spindle (MT-GFP (cyan)) relative to the mother nucleus (histone H1-RFP) within appressoria of wild-type, MoKin14 OE, and MoKin5 OE strains. All micrographs are single informative focal planes. Scale bar is 2 µm. (A) In wild-type, CKF3578, the spindle bisects the nucleus within the appressorium. (Top panel) Filled arrowheads indicate the ends of the spindle. (Bottom panel) Chromatids move asynchronously towards the ends of the spindle. (B) In the MoKin14 OE strain, CKF4106, a monopolar spindle forms within the appressorium. In both panels a relatively bright focus of MT-GFP likely represent unseparated spindle pole bodies (filled arrowhead). MTs emanate from this bright MT-GFP focus (asterisks indicate prominent MTs emanating from the unseparated SPBs). (Bottom panel) The nucleus adopts a butterfly shape, suggesting a mitotic arrest. (C) In the MoKin5 OE strain, CKF4108, a typical bipolar does not form. The spindle does not span the entire diameter of the nucleus (spindle ends marked by filled arrowheads), and the nuclear fragmentation process appears to be beginning at one end of the nucleus where the spindle is located (arrow). (D) A schematic representation of the spindle and nuclear dynamics within the appressorium corresponding to data presented in Figs. 9A-C. When MoKin14 is overexpressed, the monopolar spindle persists over time.

**Fig. 10.**
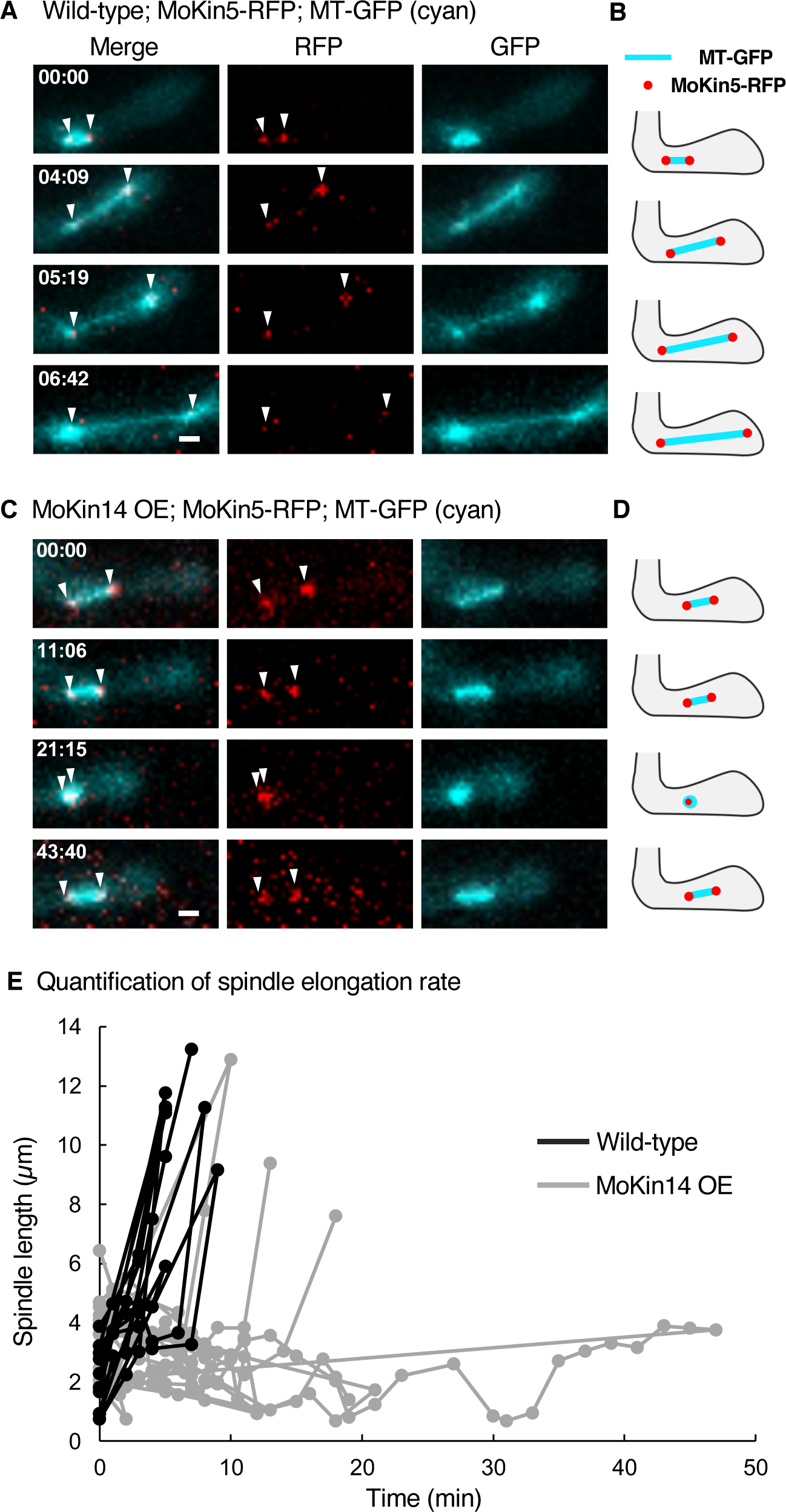
MoKin14 OE causes the spindle (MT-GFP (cyan)) to collapse and form monopolar spindles. MoKin5-RFP accumulates at the spindle pole bodies (arrowheads). All micrographs are maximum intensity projections of informative single focal planes. Scale bars are 2 µm. Time is in minutes: seconds. (A) Representative time-lapse of spindle and spindle pole body dynamics in a leading invasive hypha of wild-type *M. oryzae* strain, CKF4168. (B) Schematic representation of the spindle and spindle pole bodies dynamics as shown in Fig. 10A. (C) Time-lapse of spindle and spindle pole body dynamics in a leading invasive hypha of the MoKin14 OE strain, CKF4182. The spindle experiences several rounds of spindle collapse due to excessive MoKin14. (D) Schematic representation of the spindle and spindle pole bodies dynamics as shown in Fig. 10C. (E) Quantification of spindle length over time in invasive hyphae of the wild-type strain (CKF4168, n = 9) and the MoKin14 OE strain (CKF4182, n=24). Spindle length was determined by measuring the distance between MoKin5-tdTomato focis at the SPBs. Total time is calculated from the time the first image was acquired. The cell cycle was not synchronized, thus the spindle is not at the same spindle length at the 00:00 timepoint for each time-lapse series.

**Fig. 11.**
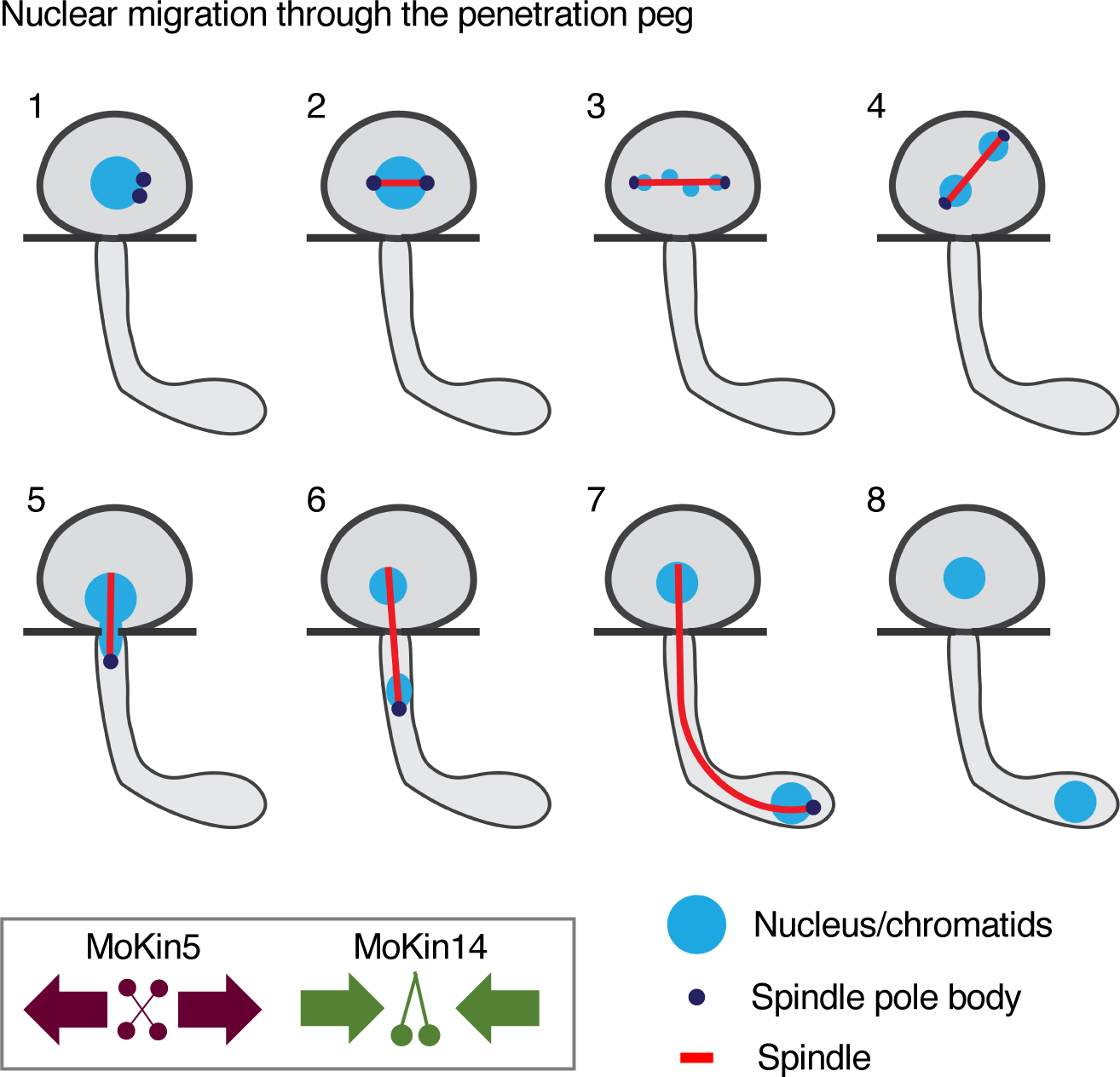
Proposed model of nuclear migration through the penetration peg. The early phases of mitosis occur within the appressorium (1–3). (1) In prophase, duplicated spindle pole bodies begin separation. (2) A bipolar spindle bisects the mother nucleus located in the appressorium in prometaphase. (3) Chromatids move asynchronously towards the spindle pole bodies in metaphase/anaphase A. (4) The spindle and divided chromatids located at the spindle pole bodies rotate to become aligned to the axis of the penetration peg in anaphase A/B. (5) The daughter nucleus begins transiting the penetration peg with the daughter bound spindle pole body leading in anaphase B. The daughter nucleus becomes highly elongated during this event. (6–7) The spindle continues to elongate, propelling the daughter nucleus towards the apical region of the primary hypha in anaphase B. (8) The daughter nucleus is positioned at the tip of the primary hypha and the spindle has collapsed, indicating exit from mitosis. The inset shows MoKin5 generating an outward force on the spindle. MoKin5 also acts as a promoter of MT nucleation. MoKin14 generates an inward force on the spindle primarily during early mitosis.

Monopolar spindles can form in two conditions. The first condition is when duplicated SPBs fail to initially separate at mitotic onset. The second condition in when duplicated SPBs fail to maintain their placement at opposite ends of the spindle throughout mitosis. In order to determine the effect of MoKin14 OE on SPBs directly, we analyzed an additional MoKin14 OE strain. This strain contained three constructs: *β-tubulin*-GFP to label the spindle; *Bas4p-MoKin14*; and *MoKin5*-RFP driven off the native *MoKin5* promoter. This particular strain was unique compared to other MoKin14 OE strains because it developed IH within the first-invaded rice cell. In both wild-type and this MoKin14 OE strain, MoKin5-RFP accumulated at the SPBs during mitosis (Fig. 10A, 10C, arrowheads). We followed the dynamics of MT-GFP and MoKin5-RFP, over time within IH of the MoKin14 OE strain, and observed a captivating pattern. The spindle experienced cycles of elongation and contraction relative to the wild-type (Fig. 10A-10D). These spindle collapse events tended to occur more frequently when the spindle was less than ∼5 µm (Fig. 10E). Yet the SPBs rapidly separated at spindle lengths exceeding ∼5 µm (Fig. 10E). We concluded that MoKin14 OE induces monopolar spindle formation due to excessive inward forces acting upon duplicated SPBs, primarily in early mitosis when the spindle is at a shorter length. The excessive inward force generated by MoKin14 OE prevented formation and maintenance of a typical bipolar spindle.

## Discussion

In this study, we demonstrated that the spindle was involved in nuclear migration through the penetration peg by genetically perturbing spindle function using an inducible overexpression promoter. We characterized the effects of kinesin-5 and kinesin-14 overexpression upon nuclear positioning, fungal development, and spindle function. Our results shed light on mechanisms permitting successful nuclear migration through the penetration peg, and the roles of kinesin-5 and kinesin-14 in the rice blast fungus, *M. oryzae.* In the following section, we discuss the mechanisms that permit nuclear migration through the penetration peg.

### Mechanisms permitting nuclear migration through the penetration peg

Our results revealed that nuclear migration through the penetration peg is initiated at the onset of mitosis within the appressorium. We observed chromatids moving towards the polar ends of the spindle in an asynchronous manner, consistent with previous studies (31–33). In our study, the spindle rotated to become aligned for movement through the appressorial pore and penetration peg. We did not observe astral MTs emanating from the spindle within the appressorium or penetration peg, although we cannot rule out that astral MTs were present but not detectable. We found live-cell imaging within the appressorium to present unique challenges in terms of visualizing fluorescently-tagged proteins that are clearly visible in other cell types of *M. oryzae.* The challenge in visualizing these fusion proteins is likely due to the highly-melanized nature of the appressorium (34). Nonetheless, our data clearly demonstrated that during nuclear migration through the penetration peg, the SPB bound to the migrating daughter nucleus proceeds the nucleus and the spindle through the penetration peg. The dynamics of the nucleus, spindle, and SPBs are summarized in Fig. 11. In migrating myoblasts from mice, the positioning of centromeres, a type of microtubule organizing centers like SPBs, is critical for effective nuclear movement (35). Interestingly, one consequence of MoKin5 OE was disruption to the arrangement of the DNA (nuclear fragments) in relation to the spindle, likely altering the position of the daughter bound SPB. The time required for the spindle to navigate towards the penetration peg was drastically increased in the MoKin5 OE strain relative to wild-type.

From these data, we propose that the daughter bound SPB plays an important role in guiding the spindle to the appressorial pore for subsequent movement through the penetration peg. The daughter bound SPB may display an enrichment of polarity determinants that help guide the spindle to the penetration peg. Similarly, the daughter bound SPB could be enriched in motor proteins, such as dynein or MoKin5, that may generate forces needed to propel the nucleus through the penetration peg. In yeast, dynein is asymmetrically distributed to one SPB, and this asymmetry is required for dynein-dependent spindle positioning at the bud neck (36). In *M. oryzae*, MoTea1 is associated with the septin and F-actin ring present near the appressorial pore where the penetration peg emerges (18). The spindle may be connecting to other cytoskeletons present at the appressorial pore via a Tea1-like mechanism as occurs in *S. pombe* (37). In *Ustilago maydis*, nuclear division defects were found in Tea1 knockout mutants in the yeast-like cells (38). More research is needed to elucidate the mechanisms that guide the daughter bound SPB efficiently to the appressorial pore in *M oryzae*.

The stretched daughter nucleus observed during movement through the penetration peg in this and a previous study is highly intriguing (23). We interpret this nuclear morphology to represent the movement of individual chromatids or clusters of chromatids through the narrow penetration peg. Recent studies show that heterochromatin levels influence nuclear migration through constricted spaces (39). One advantage of undergoing a mitotic nuclear migration through the penetration peg could be that DNA is already highly compacted into chromatids. This would allow efficient and protected movement of the nucleus through a constricted space. In *M. oryzae*, the daughter nucleus expanded in diameter immediately following movement through the penetration peg. This suggests that regions of heterochromatin within the migrating daughter nucleus relax following transit through the penetration peg. In the future, experiments altering DNA condensation within the migrating daughter nucleus may offer insight into the role DNA condensation plays in extreme nuclear migration events in *M. oryzae*.

In migratory cancer and immune cells, nuclei moving through constricted 3D spaces during interphase rely upon DNA and nuclear envelope repair mechanisms for survival (40, 41). In *M. oryzae*, which uses an intermediate form of mitosis, the outer nuclear envelope and core nucleoporins remain intact during appressorium development (42, 43), yet the behavior of the inner nuclear membrane remains undetermined. It could be that the inner nuclear membrane remains intact during nuclear migration through the penetration peg as a means to protect the nucleus. Moreover, our results suggest that nuclear migration through the penetration peg occurs during the later stages of mitosis. In other eukaryotes, ESCRT (endosomal sorting complexes required for transport) machinery is known to remodel the nuclear envelope at the later stages of mitosis (44, 45). Snf7, a component of the ESCRT-III complex, was implicated in *M. oryzae* pathogenicity on rice (46). Yet the localization of Snf7 during nuclear migration is not yet characterized. Undergoing mitosis during extreme nuclear migration through the penetration peg may allow transient nuclear envelope ruptures to be rapidly repaired by the ESCRT machinery already mobilized for mitotic function in *M. oryzae*.

### The roles of kinesin-5 and kinesin-14 in *M. oryzae*

While we provide *in vivo* evidence of MoKin5 and MoKin14 function within the spindle in *M. oryzae* during extreme nuclear migration through the penetration peg, we lack *in vitro* data to make definitive claims of the directionality of these motor proteins along MTs. In the future, *in vitro* experiments coupled with knockout experiments of MoKin5 and MoKin14 will fully elucidate whether the force-balance model of bipolar spindle formation applies to *M. oryzae*. Nonetheless, our results do provide information about the function of kinesin-5 and kinesin-14 in *M. oryzae* spindle formation and function during nuclear migration through the penetration peg. We begin this section of our discussion examining the likely roles of MoKin5 in *M. oryzae*.

#### Kinesin-5 in M. oryzae

We propose that when MoKin5 is highly overexpressed by the *Bas4* promoter, it can no longer be efficiently regulated during mitosis. This lack of regulation appears to promote excessive polymerization of MTs and excessive outward force generation, which leads to the formation of nuclear fragments. Within the appressoria of the MoKin5 OE strain, the length of the spindle continually increased in length over time. This finding is consistent with other studies of kinesin-5 overexpression. For example, overexpression of Cin8, one of two kinesin-5 motor proteins present in *S. cerevisiae*, resulted in extended spindles (47) as did kinesin-5 overexpression in the spindles of *Drosophila* embryos (48). In mice, kinesin-5 overexpression caused the formation of multipolar and monopolar spindles, and kinesin-5 overexpression was associated with polyploidy (49). In our study, MoKin5 OE caused formation of an anucleate appressorium with a single enlarged nucleus within the primary hypha. We believe it is likely this single enlarged nucleus represents a polyploid state. In mice, it is proposed that kinesin-5 overexpression prevented attachments of the chromatids to the spindle due to the generation of excessive outward forces (49). In our study, the nucleus and nuclear fragments appeared to be attached to the spindle, evident in the movement of nuclear fragments along the MT protrusions. We, therefore, favor a different mechanistic model to explain how aberrant nuclear phenotypes, and possible polyploidy, arises in the MoKin5 OE strain.

We favor a model that in the MoKin5 OE strains SPBs fail to separate. This failure in SPB separation coupled with excessive MT polymerization and excessive outward force causes the dramatic spindle and nuclear phenotypes observed in the MoKin5 OE strains. Key data from the early stages of mitosis in the appressoria support this model. We observed that a typical spindle fails to form within the MoKin5 OE strain. This was evident when a bar of MT-GFP signal spanned only approximately half the mother nucleus, and in the formation of single, double, and three or more MT protrusions. Moreover, we observed that the MoKin5 OE spindle elongates from a single plus end. If the MoKin5 OE spindle were a bipolar spindle, with each SPB maintained at the opposite end of the spindle, we would anticipate nearly equal growth from both ends of the spindle. An additional prediction is that if the MoKin5 OE spindle was a true bipolar spindle, the chromatids would move towards both poles of the spindle. This is not the pattern we found. In the MoKin5 OE strain, nuclear fragments formed along the MT protrusions, and the fragments only moved towards the growing plus-end of the spindle. We speculate that the excessive polymerization of MTs in the MoKin5 OE strain causes kinetochores to become precociously attached to MTs within the spindle. The combination of excessive MT polymerization and outward force generation causes formation of nuclear fragments. Over time, the disrupted polarity of the spindle and excessive MoKin5 causes the entire nucleus and nuclear fragments to migrate to the primary hypha.

In sum, we conclude that MoKin5 in *M. oryzae* likely generates an outward pushing force upon the spindle. We also provide data that excessive MoKin5 causes consistent polymerization of MTs within the spindle. MoKin5 OE induced distinct defects in nuclear morphology and positioning, and in spindle function compared to MoKin14 OE. We discuss the likely role of MoKin14 in the following section.

#### Kinesin-14 in M. oryzae

Our study revealed that when MoKin14 is overexpressed, spindles fail to form and maintain bipolarity throughout the early stages of mitosis. In *Aspergillus nidulans*, kinesin-14 overexpression prevents nuclear division and causes formation of monopolar spindles, consistent with our results (50). Overexpression of kinesin-14 proteins in *S. pombe* causes formation of monopolar spindles (51, 52), and overexpression of kinesin-14 in *S. cerevisiae* leads to shorter spindles (47). Given that MoKin14 OE resulted in similar spindle phenotypes, it is likely that MoKin14 generates an inward force that acts upon duplicated SPBs in early mitosis in the appressorium and in IH. However, the later stages of mitosis were relatively unaffected by MoKin14 OE. This suggests that other motor proteins, such as dynein, may be generating the antagonizing force needed to maintain the spindle in the later stages of mitosis in *M. oryzae*. There is some evidence that supports this idea. While the function of dynein in *M. oryzae* is not yet determined, knocking out a homolog of Num1, the cortical anchor of dynein, impairs nuclear positioning in vegetative hyphae, conidia, and appressoria (53). Conducting a functional study of dynein is an important future direction towards illuminating further details of nuclear migration within *M. oryzae*.

## Conclusion

The major contribution of this study is the direct evidence that the mitotic spindle mediates nuclear migration through the penetration peg in the blast fungus during colonization of the host rice cell. This knowledge is important because this is a critical step in the successful colonization of the fungus within rice tissue. Previously, the dynamics of the spindle were reported in *M. oryzae* during vegetative growth, appressorium development, IH growth, and during cell-to-cell movement through the IH peg (26, 31–33, 42, 54, 55). From these studies, we can see that delivery of a single daughter nucleus into incipient cells involves the spindle and likely occurs during the later stages of mitosis. While this finding may appear intuitive, not all fungi sync nuclear migration to nuclear division (5). Defining the contribution of the spindle during nuclear migration through the penetration peg provides fundamental knowledge about the biology of the rice blast fungus, establishes new avenues for research, and provides insight that could be exploited in the development of new anti-fungal strategies to combat blast disease.

## Materials and Methods

### Fungal and rice strains

Transgenic *M. oryzae* strains were generated by transforming wild-type O-137 (CKF558) using *Agrobacterium*-mediated transformation (56). Fungal transformants were selected on media containing either: 200 µg/mL Hygromycin (Hyg. HygR), 800 µg/mL G418 Sulfate (G418, NTPIIR), 400 µg/mL Nourseothricin (NTC, Nat1R), and 200 µM of cefotaxime (bactericide for *Agrobacterium*). Transformants were purified by single spore isolation and two to twelve independent transformants were analyzed per gene. A summary of the fungal strains, primers, constructs, and unique PCR fragments used in this study are provided in Supplemental Tables 1-4. Fungal strains were stored at -20 °C and propagated on either oatmeal agar or tomato juice agar, using standard techniques, at 24 °C with continuous light. Rice (*Oryza sativa*) cultivar YT16 was grown in a Conviron PGW36 growth chamber with daytime temperature of 28°C and nighttime temperature of 24 °C under long day conditions (14 hours/day, 10 hours/night).

### RNA isolation and gene expression analysis

The expression of *MoKin5* and *MoKin14* relative to *actin* was determined in mycelia and infected YT16 rice sheaths in two independent reverse transcription quantitative (RT-q) PCRs of wild-type (CKF3578), MoKin5 OE (CKF4108), and MoKin14 OE (CKF4106) strains. Fungal mycelia were grown in 1% sucrose complete media at 25°C for five days in a dark environment, snap frozen in liquid nitrogen, and stored at -80°C until RNA extraction. Twenty infected rice sheaths for each biological replicate (n=60 sheaths per fungal strain) were hand-trimmed and snap frozen in liquid nitrogen at 30-31 hours post inoculation. We confirmed that each fungal strain had penetrated into the rice tissue by conducting confocal microscopy two hours prior to harvesting the sheath samples (data not shown). For mycelia and infected sheath samples, total RNAs were extracted using the Trizol method combined with the RNA Clean and Concentrator -5 kit (Zymo), according to manufacturer’s instructions. Genomic DNA was removed using Turbo™ DNase (Ambion) using manufacturer’s instructions. Complementary DNA (cDNA) was synthesized following manufacturer’s instructions using the ImProm II Reverse Transcriptase system (Promega) from 500 ng of total RNAs for mycelial samples and 650 ng of total RNAs for sheath samples. Applied Biosystems SYBR Green qPCR 2X Master Mix (Thermo Fisher) was used to perform the RT-qPCRs with a CFX96 Touch Real-Time PCR Detection System (BioRad). Reactions contained 7 µL Applied Biosystems SYBR Green qPCR Master Mix, 1.5 µL each of the forward and reverse primer (3.3 nM concentration, Table S2), 1.5 µl cDNA, and 2.5 µL distilled water, for a final volume of 14 µL. Standard thermocycling conditions for primers ≥ 60°C per the Applied Biosystems SYBR Green qPCR Master Mix manufacturer’s instructions were used. Thermocycler conditions were: 2 minutes at 50°C, 2 minutes at 95°C, and 40 cycles of 15 seconds at 95°C and 1 minute at 60°C. Relative expression levels of *MoKin5* and *MoKin14* were calculated using the *M. oryzae actin* gene (MGG_03982) as reference (57). The 2-^ΔΔCt^ was used to calculate relative expression levels (58). Average threshold cycle (Ct) values from three technical replicates were normalized to *actin* for each strain (ΔCt). This value was subtracted from the calculated mean ΔCt value of the wild-type (CKF3578) in the respective mycelia or sheath condition, yielding the ΔΔCt value. These values were transformed using the equation 2-^ΔΔ^Ct. Mean 2-ΔΔCt values, along with 95% confidence intervals, were calculated for each strain from three biological replicates.

### Pathogenicity assays and live-cell imaging

#### Rice sheath inoculations

Susceptible rice cultivar YT16 was inoculated with fungal spores as described previously (59). Leaf sheaths 3-8 cm in length from 2 to 3-week old plants were inoculated with either 3-4 X 104 spores per mL for ∼48 hour post inoculation (hpi) observation, or 7-10 × 10^4^ spores per mL for ∼28 hpi observation. All spore inoculum was filtered using Miracloth. Inoculated sheaths were prepared for microscopy by hand trimming with razor blades.

#### Appressorium development assay

Spores were harvested and diluted to a final concentration of 2-4×10^4^. Spores were inoculated onto a hydrophobic coverslip and incubated for 3-4 hours at room temperature prior to microscopy.

#### Whole-plant spray inoculations

Spores were collected from 7 to 10-day old V8 tomato juice agar plates and diluted to a final concentration of 1×10^5^ spores per mL in 0.2% gelatin. 17-day old YT16 rice plants were sprayed with 5 mL of spores. Sprayed rice plants were placed in clear plastic bags overnight at room temperature. The next day sprayed plants were removed from the plastic bags and placed in a Conviron PGW36 growth chamber with daytime temperature of 28°C and nighttime temperature of 24 °C under long day conditions. Infected leaves were harvested 7 days after inoculation. Infected leaves were harvested and collected on notecards that were scanned with an Epson Perfection 4870 Photo Scanner to generate digital images for analysis of lesion development. Lesion development was analyzed using ImageJ to determine the percentage of diseased tissue area. Briefly, scanned leaf images were color adjusted to find the total area of the leaf and then again adjusted to find the total area of the diseased tissue. The resulting ratio was converted to a percentage for each biological replicate, and mean values along with margins of error were calculated for each fungal strain. Figures of infected rice leaves were compiled with Adobe Photoshop and Adobe Illustrator.

#### Confocal microscopy and analysis

Live-cell confocal microscopy of developing appressoria and infected rice sheaths was conducted using a Zeiss 880 confocal system equipped with a Plan-Neofluor 40×/1.3 NA (oil) objective. Excitation/emission wavelengths were 488 nm/505–530 nm (GFP), and 543 nm/560– 615 nm (RFP). Analyses of resulting micrographs were done using combinations of the Zen software (Black and Blue editions). Figures were compiled using Zen software (Black and Blue editions), Adobe Illustrator, and Microsoft PowerPoint.

#### Quantification of nuclear phenotypes

Informative micrographs collected from wild-type, MoKin5 overexpression (OE), and MoKin14 OE strains at approximately 28 hours post inoculation and 48 hours post inoculation were analyzed. Only infection sites with an intact appressorium and developed primary hypha were considered for quantification. Observed patterns of nuclear positioning within the appressorium and primary hypha were quantified. Phenotype frequency was compiled and graphed using a combination of Microsoft Excel and Adobe Illustrator.

#### Quantification of rate of mitosis

Rate of mitosis in wild-type and MoKin14 OE strains was determined using the time the first micrograph in a time-lapse series was acquired as the 00:00 timepoint. The spindle length was calculated by selecting a single informative focal plane, and measuring the length of the spindle from SPB to SPB (marked by MoKin5-RFP) using the line tool in Zen Black. In monopolar spindles, the length of spindle was measured using the MT-GFP fluorescence signal. The resulting time and spindle length intervals were analyzed and plotted in Microsoft Excel.

#### Quantification of spindle length in the MoKin5 OE strain

The length of the spindles observed in strain CKF4203 was determined using the Closed Bezier tool in Zen software (Black edition). The length of the spindle was measured from the minus-end to the plus-end.

### Sequence information

Gene identification numbers, except for *actin* and *Bas4*, were determined using *Aspergillus nidulans* or *Schizosaccharomyces pombe* protein sequences as query sequences in NCBI BlastP searches of the non-redundant protein sequence database using the *Magnaporthe oryzae 70-15* reference genome. Protein sequences were obtained from FungiDB. Gene identification numbers for *M. oryzae* were identified and gene sequence information along with 2 Kb upstream and downstream was downladed from FungiDB and analyzed using Geneious Prime 2019.2.3. Reciprocal NCBI BlastP searches using the protein and gene sequences from *M. oryzae* to either *A. nidulans* or *S. pombe* were conducted as a quality control step. A list of resulting gene identification numbers is available in Supplemental Table 2. Sequence information is available from FungiDB.

### Statistical analysis and reproducibility

Significance of gene expression levels was determined using a Student’s two-tailed test assuming unequal variance in Microsoft Excel. Significance of nuclear positioning phenotypes at the early timepoint was determined using Fisher’s exact test (60) in in GraphPad QuickCalcs (accessed March 19, 2021). Confocal micrographs are representative of at least three biological replicates. Representative examples of each strain are presented throughout the figures.

## Data availability

Data supporting the conclusions of this study are available upon reasonable request from the corresponding author. Key constructs generated in this study will be made available from Addgene. Mutant fungal strains are available from corresponding author with appropriate permits.

## Acknowledgements

We thank current and past members of the Khang Lab for their support and feedback, especially Kathy Prado, Brandon Mangum, Jie Zhu, Kiersun Jones, and Dong Won Kim. We are grateful to the Devos lab at the University of Georgia for access to the equipment needed to complete the gene expression analyses. We also thank the University of Georgia Fungal Group for iterative feedback over the lifetime of the project. We acknowledge the assistance of the Biomedical Microscopy Core at the University of Georgia with imaging using a Zeiss LSM 880 confocal microscope. This work was supported by the National Science Foundation BREAD program under Grant No. 1543901 (CHK) and the National Science Foundation Graduate Research Fellowship Program under Grant No. 1842396 (MAP). Any opinions, findings, and conclusions or recommendations expressed in this material are those of the author(s) and do not necessarily reflect the views of the National Science Foundation.

## Supplemental Figure Legends

**Fig. S1.** MoKin5 is a conserved kinesin-5. (A) Schematic of MoKin5 protein structure. A PFAM kinesin motor domain (PF00225) is predicted at positions 109 to 438. (B) Protein sequence alignment of MoKin5 and kinesin-5 in *Aspergillus nidulans* (AnBimC; AN3363). Dashes represent indels, and dots represent mismatched amino acids. Magenta box corresponds to predicted kinesin motor domain in Fig. S1A. (C) Protein sequence alignment of MoKin5 to select kinesin-5 proteins in other fungi, including *A. nidulans* (AnBimC), *Schizosaccharomyces pombe* (SpCut7), and *Saccharomyces cerevisiae* (ScKip1, ScCin8).

**Fig. S2.** Relative localization of MoKin5-RFP and Mo-γ-tubulin-RFP in spindles (MT-GFP) spanning the germtube of developing appressoria (asterisks). Micrographs are single informative focal planes. Scale bars are 2 µm. (A) Localization of MoKin5-RFP (black arrowheads) relative to MT-GFP in a developing appressorium of *M. oryzae* strain CKF4168. Gray arrow shows red autofluorescence in micrograph. (B) Localization of Mo-γ-tubulin-RFP (black arrowheads) relative to MT-GFP in a developing appressorium of *M. oryzae* strain CKF4117. CKF4117 displayed severe developmental defects and was not used for subsequent analysis. (C) Protein sequence alignment of Mo-γ-tubulin (Mogamma; MGG_00961) to γ-tubulin in *Aspergillus nidulans* (Angamma; MipA; AN0676). Dashes represent indels, and dots represent mismatched amino acids.

**Fig. S3.** Summary of observed nuclear positioning and morphology phenotypes. Only infection sites with intact appressoria were considered for analysis. For each strain, only micrographs with histone H1-RFP and brightfield channels were scored. (A) Frequency of all nuclear positioning and morphology phenotypes at ∼28 hours post inoculation in wild-type (CKF3578, CKF3971, n=153), MoKin5 OE (CKF4108, n=125), and MoKin14 OE (CKF4106; CKF4093, n=85) strains. Schematic representations of nuclear positioning and morphology for each group are found at the bottom of Fig. S3B. (B) Frequency of all nuclear positioning and morphology phenotypes at ∼48 hours post inoculation in wild-type (CKF3971, n=26), MoKin5 OE (CKF4108, n=46), and MoKin14 OE (CKF4093, n=56) strains. Schematic representations summarize key nuclear positioning and morphologies for each group. Group 1, Group 5, and Group 7 example micrographs can be found in Fig. 5A. A Group 2 example micrograph is found in Fig. 6B. A Group 3 example micrograph is found in Fig. S3C. A Group 4 example micrograph is found in Fig. 7. A Group 6 example micrograph is found in Fig. 6C. (C) An example micrograph of a Group 3 nuclear phenotype. Single focal planes are shown. A nucleus (arrow) appears to be stuck in the penetration peg. The appressorium is indicated by an arrowhead. Scale bar is 2 µm.

**Fig. S4.** Representative examples of wild-type, MoKin5, and two independent MoKin14 OE strains in infected rice sheaths at ∼48 hours post inoculation. Micrographs are single focal planes. Scale bars are 10 µm. Arrowheads point to appressoria.

**Fig. S5.** Representative example of localization of MoKin5-RFP and MT-GFP in a MoKin5 OE strain CKF4203. Arrowheads point to appressoria. The appressorium designated with “M” is in mitosis, the spindle is evident (arrow) and MoKin5-RFP fails to localize at the ends of the spindle, as occurs in wild-type. The appressorium designated with “I” is in interphase, and MoKin5-RFP localizes within the nucleus, as occurs in wild-type. Micrograph is a single informative focal plane. Scale bar is 5 µm.

**Fig. S6.** MoKin14 is a conserved kinesin-14. (A) Schematic of MoKin14 protein structure. A PFAM kinesin motor domain (PF00225) is predicted at positions 554 to 881. (B) Protein sequence alignment of MoKin14 and kinesin-14 in *Aspergillus nidulans* (AnKlpA; AN6340). Dashes represent indels, and dots represent mismatched amino acids. Green box corresponds to predicted kinesin motor domain in Fig. S6A. (C) Protein sequence alignment of MoKin14 to select kinesin-14 proteins in other fungi including *A. nidulans* (AnKlpA), *Schizosaccharomyces pombe* (SpKlp2, ScPkl1), and *Saccharomyces cerevisiae* (ScKar3).

